# High current production of *Shewanella oneidensis* with electrospun carbon nanofiber anodes is directly linked to biofilm formation

**DOI:** 10.1101/2021.01.28.428465

**Authors:** Johannes Erben, Xinyu Wang, Sven Kerzenmacher

**Affiliations:** Center for Environmental Research and Sustainable Technology (UFT), University of Bremen, 28359 Bremen, Germany; Laboratory for MEMS Applications, IMTEK – Department of Microsystems Engineering, University of Freiburg, Georges-Koehler-Allee 103, 79110 Freiburg, Germany

**Keywords:** Biocatalysis, Bioelectrochemical systems, Electrospinning, Macropore, Material science, Morphology, *Shewanella oneidensis* MR-1, Surface area

## Abstract

We study the current production of *Shewanella oneidensis* MR-1 with electro-spun carbon fiber anode materials and analyze the effect of the anode morphology on the micro-, meso-and macroscale. The materials feature fiber diameters in the range between 108 nm and 623 nm, resulting in distinct macropore sizes and surface roughness factors. A maximum current density of (255± 71) µA cm^−2^ was obtained with 286 nm fiber diameter material. Additionally, micro-and mesoporosity were introduced by CO_2_ and steam activation which did not improve the current production significantly. The current production is directly linked to the biofilm dry mass under anaerobic and micro-aerobic conditions. Our findings suggest that either the surface area or the pore size related to the fiber diameter determines the attractiveness of the anodes as habitat and therefore biofilm formation and current production. Reanalysis of previous works supports these findings.

## 2 Introduction

### 2.1 *Shewanella oneidensis* MR-1 and its applications in Bioelectrochemical Systems

Due to recent attempts to find alternatives for the fossil-based economy, Bio-electrochemical systems (BES) moved into the focus of scientific interest. In BES, oxidative and reductive reactions are catalyzed by, e.g., electroactive bacteria. Electrodes are used to establish an electrical connection between bacteria cells and an external electrical circuit. The most prominent applications of BES comprise electricity production with microbial fuel cells [Logan et al., 2006], microbial electrosynthesis [Nevin et al., 2010] for carbon dioxide fixation, and more recently, electrode-assisted fermentation for the production of platform chemicals [Bursac et al., 2017, Flynn et al., 2010, Förster et al., 2017]. In this work, the interplay between the electroactive microbe Shewanella oneidensis MR-1 (MR-1), a naturally occurring facultative anaerobic bacterium, and electrospun carbon nanofiber electrodes is studied. MR-1 is not only a well establish model organism for microbial anodes, but due to its fully sequenced and annotated genome [Heidelberg et al., 2002] and ease of cultivation, it is a well-suited chassis organism for the genetic engineering of production strains [Bursac et al., 2017, Flynn et al., 2010]. The extracellular electron transfer (EET) to insoluble terminal electron acceptors of MR-1 is subject to extensive scientific investigation. Three distinct electron transfer mechanisms were proposed for MR-1: mediated electron transfer (MET) via soluble redox-active species, direct electron transfer (DET), and electron transport via pili [Carmona-Martinez et al., 2011]. The ability to export electrons across the cellular membrane stems from several multi-heme c-type cytochromes bridging the gap between the cytoplasm and cell surface [Okamoto et al., 2011]. It has been shown that self-secreted flavins, most of them being riboflavin (RF), increase the current production of MR-1 [Marsili et al., 2008]. The current production increase was attributed to MET via redox-cycling of RF, using knock-out mutants and medium replacement experiments [Carmona-Martinez et al., 2011, Marsili et al., 2008]. Okamoto et al. [2013, 2015] suggested a different role of flavins in the extra-cellular electron transport: flavin mononucleotide and RF act as co-factor of the outer membrane cytochromes MtrC and OmcA respectively and enhance their extracellular electron transfer rates. This role may explain the observed current increase in the presence of low concentrations of flavins that hardly can be accounted for by diffusive MET [Torres et al., 2010].

### 2.2 Biofilm formation of *Shewanella oneidensis* MR-1

The achievable current density with MR-1 strongly depends on the number of cells attached to the electrode. The operation conditions within a bioelectrochemical reactor and the electrode’s working potential affect the biofilm formation of MR-1 cells on electrodes [Carmona-Martínez et al., 2013]. The biofilm coverage on electrodes is not limited to single layers and can be stimulated by the presence of oxygen [Kitayama et al., 2017], possibly due to increased motility or motility elements that promote the development of three-dimensional biofilm structures [Thormann et al., 2004]. In static multilayer biofilms, electron transport from cells that are not directly attached to the electrode is facilitated by the sequential redox cycling of outer membrane c-type cytochromes and/or MET [Pirbadian et al., 2020]. In dynamic biofilms, motile cells could increase the apparent cell surface coverage [Okamoto et al., 2012], enabling more cells to participate in EET. Besides oxygen, RF in solution, RF covalently bound to magnetic beads, and polymerized RF on electrodes are known to stimulate biofilm growth on electrodes, and a positive correlation between biofilm formation and current production was observed [Arinda et al., 2019, Zou et al., 2019]. More positive anode working potentials have been shown to enhance biofilm formation and current output [Carmona-Martínez et al., 2013]. While increased current production is preferential for a high production rate, a lower coulombic efficiency (CE) decreases production yield.

### 2.3 Anode materials tailored for the operation with *Shewanella oneidensis* MR-1

Compared to *Geobacter sulfurreducens*, MR-1 produces lower current densities, but the current production can be improved by choosing a suitable anode material [Kipf et al., 2014]. Numerous studies with mixed-community biofilms have been conducted on anode materials [Chen et al., 2011, Flexer et al., 2013, He et al., 2011, Picot et al., 2011], while little systematic work with MR-1 is reported in the literature. Kipf et al. [2013] identified a commercial steam-activated knitted carbon fiber material with a current density of 24 µA cm^−2^ at a potential of −200 mV (sat. Calomel; 41 mVvs NHE) as superior to other carbon-based materials. Patil et al. [2013] studied the current production of electrospun carbon fiber materials partly modified with activated carbon and graphite particles. Increased current production was observed with increasing specific BET surface area. Pötschke et al. [2019] investigated the carbon composition and fiber breaks. They identified a combination of highly conductive graphitic carbon, and free filament ends as key to high current densities. In the present study, we systematically investigate the effect of the electrode the surface area, and the pore size on the anodic current production and biofilm formation of MR-1 under anaerobic and micro-aerobic conditions. The controlled fabrication process of the materials allows us to separate effects of the macro, meso-and micro-scale morphology on the current production. The surface chemistry is presumably identical, allowing us to separate the effects of the material morphology and the surface chemistry on the current production.

We tested four materials with distinct fiber diameters that feature surface roughness factors (*SRF* s) between 286 and 1004 and mean macropore diameters between 0.4 µm and 1.6 µm. Additionally, the electrospun carbon fiber surface was modified by steam and CO_2_ activation to increase the surface area further and introduce micro-and mesopores, which have been hypothesized to enhance MET [Kipf et al., 2013, Patil et al., 2013]. We also reevaluate previous systematic anode material studies [Kipf et al., 2013, Patil et al., 2013, Pötschke et al., 2019] based on the premise that mainly the macrostructure of the anode determines the current production and biofilm formation of MR-1. The commercial reference materials were investigated previously [Dolch et al., 2014, Förster et al., 2017, Kipf et al., 2013, 2014] and serve as a benchmark in this work.

## 3 Results and Discussion

### 3.1 Anode material properties

The pore sizes and surface area of the anode materials were characterized on the micro (*<*2 nm), meso (*>*2 nm to *<*50 nm), and macro (*>*50 nm) scale by nitrogen adsorption, electrochemically accessible surface (*ECAS*), and SEM imaging. Scanning electron microscopy photographs of the materials are provided in Fig. S1.1. The self-made electrospun materials ES100 to ES600 feature ultra-thin fibers with mean fiber diameters between ∼100 nm and ∼600 nm. The commercial reference materials GFD 2 and C-Tex 13 consist of PAN derived fibers with about 7 µm to 8 µm diameter. The individual activated C-Tex 13 fibers feature grooves of about 1 µm width along the fiber (see inset of Fig. S1.1 E). The fibers are loosely twisted in bundles that are knitted into a fabric. GFD 2 felt consists of mostly smooth graphite fibers. C-Tex 13 and the activated electrospun materials feature a hydrophilic surface. GFD 2 and the electrospun non-activated materials are hydrophobic. The carbonization process is the same for all electro-spun materials. The electrospun materials and GFD 2 exhibit porosities from 94 % to 95 %. C-Tex 13 has a lower porosity of 81 %. The morphological characteristics of all investigated materials are listed in Tab. 1 and are discussed in detail below.

**Table 1:**
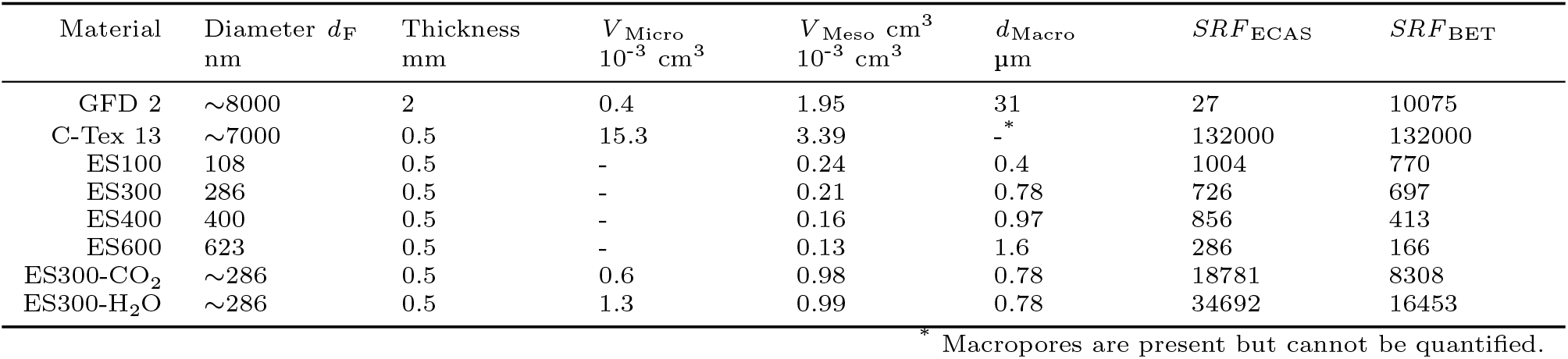
Properties of the commercial and electrospun anode materials. The complete material properties are listed in Tab. S3.1 in the supplementary material. The pore diameters are: Micro: *<* 2 nm, Meso: *>* 2 nm & *<* 50 nm, Macro: *>* 50 nm. The material thicknesses are given according to the manufacturer’s data sheet and have not been verified by own measurements, except for the electrospun materials and GFD 2.

| Material | Diameter $d_F$<br>nm | Thickness<br>mm | $V_{\text{Micro}}$<br>$10^{-3}\text{ cm}^3$ | $V_{\text{Meso}}$ $\text{cm}^3$<br>$10^{-3}\text{ cm}^3$ | $d_{\text{Macro}}$<br>$\mu\text{m}$ | $SRF_{\text{ECAS}}$ | $SRF_{\text{BET}}$ |
| --- | --- | --- | --- | --- | --- | --- | --- |
| GFD 2 | $\sim 8000$ | 2 | 0.4 | 1.95 | 31 | 27 | 10075 |
| C-Tex 13 | $\sim 7000$ | 0.5 | 15.3 | 3.39 | -* | 132000 | 132000 |
| ES100 | 108 | 0.5 | - | 0.24 | 0.4 | 1004 | 770 |
| ES300 | 286 | 0.5 | - | 0.21 | 0.78 | 726 | 697 |
| ES400 | 400 | 0.5 | - | 0.16 | 0.97 | 856 | 413 |
| ES600 | 623 | 0.5 | - | 0.13 | 1.6 | 286 | 166 |
| ES300-$\text{CO}_2$ | $\sim 286$ | 0.5 | 0.6 | 0.98 | 0.78 | 18781 | 8308 |
| ES300-$\text{H}_2\text{O}$ | $\sim 286$ | 0.5 | 1.3 | 0.99 | 0.78 | 34692 | 16453 |

\* Macropores are present but cannot be quantified.

The steam activated C-Tex 13 exhibits the highest *SRF* _ECAS_, *SRF* _BET_, and micropore volume *V* _Micro_ among the investigated materials. Interestingly, the graphitic GFD 2 also features a substantial micropore volume but a lower *SRF* _ECAS_ of 27. Graphite contains lesser oxygen-containing groups [Ko et al., 1992] that increase the double layer capacitance [Bleda-Martínez et al., 2005] and thus the *ECAS*. The high mesopore volumes *V* _Meso_ of 3.39×10^−3^ cm^3^ (C-Tex 13) and 1.95× 10^−3^ cm^3^ (GFD 2) are mainly due to the higher area densities of 182 g m^−2^ and 165 g m^−2^. The mean macropore diameter *d* _Macro_ of GFD 2, 31 µm, is about a factor of 20 higher than the electrospun materials with similar morphology. The macropore structure of C-Tex 13 can only be assessed semi-quantitatively due to its knitted structure: The grooves along the fibers introduce porosity in the submicron range, and interfiber distances vary from direct contact to several fiber diameters within one fiber bundle. Maximum distances between the fiber bundles are in the range of ∼.200 µm. The electrospun materials’ *ECAS* values do not monotonically increase with decreasing fiber diameter, possibly due to the fiber cross-section’s deviation from a perfectly round shape [Erben et al., 2021] or differing surface roughness. The same holds for the *SRF* _ECAS_ due to similar area densities. However, *S* _BET_ and *SRF* _BET_ increase monotonically with decreasing fiber diameter. CO_2_ and steam activation increase the *SRF* _ECAS_ of ES300 by 30 and 55 and *SRF* _BET_ by a factor of 12 and 24, respectively. The increased micro- and mesopore volume of the activated electrospun materials show that the increased *SRF* s are linked to micro- and mesopore formation during activation. Because of the low weight loss during activation, the fiber diameter and macropores remain practically unaffected by CO_2_ and steam activation.

The steam activated C-Tex 13 exhibits the highest *SRF* _ECAS_, *SRF* _BET_, and micropore volume *V* _Micro_ among the investigated materials. Interestingly, the graphitic GFD 2 also features a substantial micropore volume but a lower *SRF* _ECAS_ of 27. Graphite contains lesser oxygen-containing groups [Ko et al., 1992] that increase the double layer capacitance [Bleda-Martínez et al., 2005] and thus the *ECAS*. The high mesopore volumes *V* _Meso_ of 3.39×10^−3^ cm^3^ (C-Tex 13) and 1.95 ×10^−3^ cm^3^ (GFD 2) are mainly due to the higher area densities of 182 g m^−2^ and 165 g m^−2^. The mean macropore diameter *d* _Macro_ of GFD 2, 31 µm, is about a factor of 20 higher than the electrospun materials with similar morphology. The macropore structure of C-Tex 13 can only be assessed semi-quantitatively due to its knitted structure: The grooves along the fibers introduce porosity in the submicron range, and interfiber distances vary from direct contact to several fiber diameters within one fiber bundle. Maximum distances between the fiber bundles are in the range of ∼200 µm. The electrospun materials’ *ECAS* values do not monotonically increase with decreasing fiber diameter, possibly due to the fiber cross-section’s deviation from a perfectly round shape [Erben et al., 2021] or differing surface roughness. The same holds for the *SRF* _ECAS_ due to similar area densities. However, *S* _BET_ and *SRF* _BET_ in-crease monotonically with decreasing fiber diameter. CO_2_ and steam activation increase the *SRF* _ECAS_ of ES300 by 30 and 55 and *SRF* _BET_ by a factor of 12 and 24, respectively. The increased micro- and mesopore volume of the activated electrospun materials show that the increased *SRF* s are linked to micro- and mesopore formation during activation. Because of the low weight loss during activation, the fiber diameter and macropores remain practically unaffected by CO_2_ and steam activation.

### 3.2 Electrical resistivity of the anode materials

The sheet resistances for the electrospun materials given in Erben et al. [2021] allow the estimation of the ohmic potential drop across the electrode according to Madjarov et al. [2017]. For the electrospun materials and GFD 2, with sheet resistances between ∼1.8 Ω and ∼2 Ω, the ohmic drop for the current densities of MR-1 obtained in this work is estimated to be less than 1 mV. C-Tex 13 has a higher sheet resistance of 144 Ω, resulting in an ohmic drop of less than 40 mV at the maximum current densities measured with the materials. Therefore, we do not expect that the different electrical resistances affect the current production.

### 3.3 Current production under anaerobic and micro-aerobic conditions

The six different anode materials were characterized in four technical replicates to account for the experimental conditions’ variability. The values reported in this section are averages with sample standard deviation (n = 4). As one anode material of a kind was used for biofilm imaging by SEM, three values were considered in the biofilm dry weight analysis in Section 3.4. The maximum current densities presented in Fig. 1 A were extracted from chronoam-perometry data recorded over the experimental period of 2 weeks (data not shown). Under anaerobic conditions (Fig. 1A, red), a maximum current density of (255±71) µA cm^−2^ was recorded with the ES300 material. This current density is about 1.4-fold higher than the highest value of 186 µA cm^−2^ reported up to now Pötschke et al. [2019]. A trend towards higher current densities with decreasing fiber diameter is apparent (Fig. 1 A, red). Through the fiber diameter *d* _F_ and the porosity *φ, SRF* and macropore diameter *d* _Macro_ are inseparably linked in fibrous materials on the macroscale (a detailed derivation of the analytical model can be found in Section S4.1 of the supplementary information):

**Figure 1:**
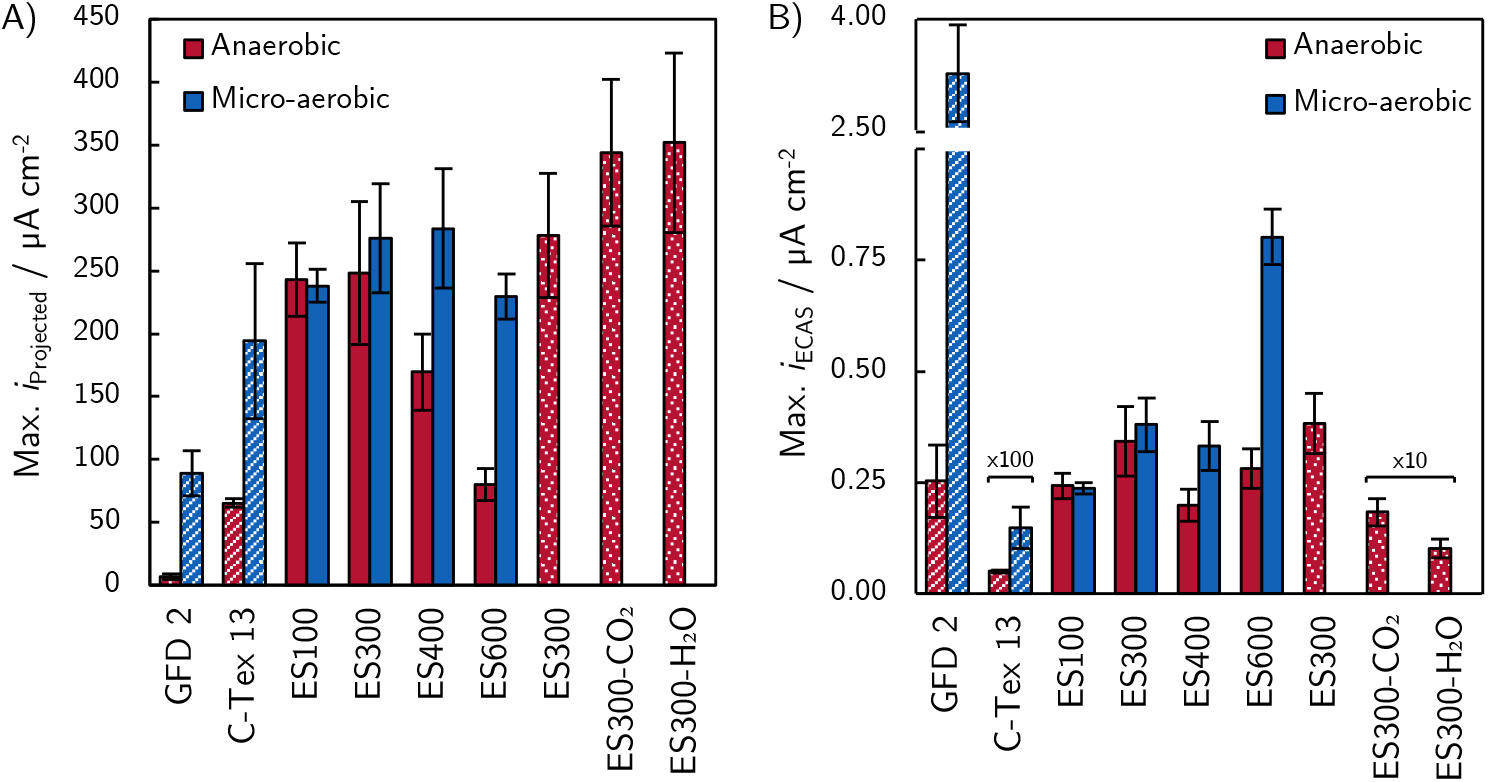
A) The maximum current normalized to the projected surface area. The hatched filling indicates the commercial reference materials, and the dotted filling the additional technical and biological replicates with activated electrospun materials conducted in a separate set of experiments. The values for C-Tex 13 were multiplied by 100 and the values of ES300-CO_2_ and ES300-H_2_O by a factor of 10 for better visualization. All values represent arithmetic means and sample standard deviation and are calculated based on a quadruplicate. A) A trend to higher current densities with decreasing fiber diameter is apparent under anaerobic conditions. Welch corrected t-tests did not reveal a significant improvement of the current production of activated ES300 materials ES300-CO_2_ (*p* = .14, Welch corrected t-test) and ES300-H_2_O (*p* = .15) compared to the non-activated ES300 material (Fig. S2.5). Under micro-aerobic conditions, similar current productions of the electrospun materials and C-Tex 13 are observed. The current production of GFD 2 is increased 13-fold under micro-aerobic conditions compared to anaerobic conditions. B) The maximum current density normalized to *ECAS* reveals that the current production is not directly linked to the internal surface area.

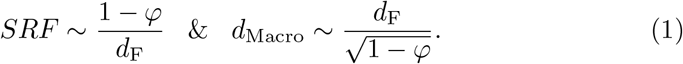

As predicted by the analytical model, *SRF* _ECAS_, *SRF* _BET_, and *d* _Macro_ intercor-relate strongly for the non-activated electrospun materials (absolute correlation coefficients between 0.82 and 0.96, Tab. S3.2). Additionally, we find that the mesopores volume *V* _Meso_ is correlated to the *SRF* and *d* _Macro_ (absolute correlation coefficients between 0.79 and 0.96). As expected, the maximum current density can be correlated reasonably well with all of the above material properties (Fig. S2.4). Therefore, the increased current production of thin fiber material cannot be attributed to the increased *SRF*, reduced macropore diameter, or the increasing mesopore volume. Steam and CO_2_ activation introduce micro and mesopores, while the macropore structure remains unchanged. The surface roughness factor increases due to the surface within the micro and mesopores, but the additional surface does not significantly improve current production (Fig. S2.5). This allows us to rule out micro- and mesopore’s positive effects on the current production and the related pore surface. We will further discuss the role of macropore size and *SRF* for the current production again in Section 3.7. Under micro-aerobic conditions (Fig. 1 A, blue), the different non-activated electrospun anode materials show maximum current densities between (234±16) µA cm^−2^ and (305± 58) µA cm^−2^. The presence of trace amounts of oxygen under micro-aerobic conditions masks thin fibers’ positive effect on the current production. Due to the strong variability of the obtained values, no significant difference in the current production can be observed. Although oxygen serves as a competing electron acceptor to the anode, it enhances current production [Kipf et al., 2013] and bacterial growth and biofilm formation (see Section 3.4). Little systematic research on the influence of the anode macro morphology on the current production has been conducted up to now. Diverse characterization techniques make it impossible to compare results directly between different studies. The current production of the commercial reference materials C-Tex 13 and GFD 2 was characterized previously using a quasi-galvanostatic protocol by Kipf et al. [2013]. Analysis of the experimental setup in retrospect revealed that trace amounts of oxygen were present in the reactor. Therefore, micro-aerobic conditions in this work are comparable to the anaerobic conditions in Kipf et al. [2013]. Interestingly, the commercial reference materials C-Tex 13 and GFD 2 exhibit about 10-fold higher current densities, with (229± 82) µA cm^−2^ and (112± 36) µA cm^−2^ respectively, than previously reported [Kipf et al., 2013]. The 10-fold higher current production is most likely a result of the different characterization methods. In the present study, even e.g. trace amounts of oxygen change the difference in the reference materials’ current production from 9-fold under micro-aerobic to 2-fold under micro-aerobic conditions. This underlines the necessity of a systematic and well-controlled approach to material characterization. In the following, we reevaluate data from previous publications based on the premise that the anode materials’ macrostructure determines its current production with MR-1 taking into account the possible effects of oxygen contamination. Tab. S5.3 summarizes the data of Kipf et al. [2013] and includes additional morphological characteristics. We find that knitted, and woven activated carbon fabrics’ current production is significantly higher than the carbon felts and carbon paper (*p* ¡ .001, Welch corrected *t*-test). The current production of felt materials, made of grooved activated carbon fibers (V1-Felt) and smooth graphitized fibers (GFD 2), does not differ significantly (*p* = .13). This confirms our finding that micro- and mesopores do not enhance the current production considerably. Within the group of knitted and woven fabric anode materials, C-Tex 27 produces a significantly lower current than C-Tex 13 (*p* = .0018). Judging from the SEM images, the densely aligned fibers of C-Tex 27 and woven structure cause less accessible pore space in and between the fiber bundles compared to C-Tex 13, consisting of knitted twisted fiber bundles. C-Tex 20 features a higher area density and thickness than C-Tex 13, while the fiber diameter is the same. The higher current production of C-Tex 20 compared to C-Tex 13 (*p* = .069) can therefore be attributed to the increased area-specific volume due to the higher thickness of the electrode material, similar to four layers of C-Tex 13 compared to a single layer [Kipf et al., 2013]. It is noteworthy that the current production does not scale linearly with the number of layers, possibly to mass transport limitations. Patil et al. [2013] investigated the current production of electro-spun PAN-based carbon fibers without (PAN) and with additional activated carbon (PAN-AC) and graphite (PAN-GR) particles (SEM images and material properties see Tab. S5.4). The higher current production of the materials with particles was attributed to their higher surface area. Besides higher BET surface area and micropore volume of PAN-AC and PAN-GR, the materials with particles and a higher surface area also exhibited smaller fiber diameters. Similar to our results, the increasing current production with decreasing fiber diameter could result from the macrostructure with a larger accessible surface area and smaller macropores. In a study by P ö tschke et al. [2019], the effect of the chemical carbon fiber composition and fiber breaks on the current production was investigated. SEM images of fiber bundles depicted in Tab. S5.5 reveal that the fibers of stretch broken (SB) fabrics are less aligned than non-broken (CM) fibers with pores between the individual fibers. However, the mean current production of SB fabrics compared to CM fabrics is not significantly higher (*p* = .32). However, planktonic cell growth (max. *OD* _600_ ∼0.4 to ∼0.9) and flavin accumulation (∼0.4 µM to ∼0.8 µM) are indications of trace amounts of oxygen present in this study (see Section 3.6) that mask effects of the anode morphology on the current production. Besides oxygen, the higher anode potential of 200 mV (sat. Ag/AgCl, 400 mV vs. NHE) compared to the 0 mV (sat. Ag/AgCl, 200 mV mV vs. NHE) in our study might enable cell growth and flavin secretion.

### 3.4 Current production and biofilm formation

The bacterial dry weight was quantified after two weeks of operation (Fig. 2 A). Under micro-aerobic conditions, a higher bacterial dry mass was found for all anode materials compared to anaerobic conditions. Under micro-aerobic conditions, similar amounts of bacterial dry mass with large variability between 38 % and 73 % were found on the electrospun materials and the commercial reference materials. Under anaerobic conditions, however, increasing bacterial dry mass with decreasing fiber diameter can be observed for the electrospun materials (Fig. 2 A, plain red). The similarity to the maximum current densities in Fig. 1 A is apparent. Therefore, the final current densities (after two weeks of operation) were normalized to the bacterial dry weight (Fig. 2 B). Most interestingly, the electrospun materials show similar values as would be expected for a proportional relationship between the number of current producing cells and current production for anaerobic and micro-aerobic conditions. The values of C-Tex 13 under both operating conditions, and GFD 2 under micro-aerobic conditions, do not deviate significantly from the average value of (65.2±7.0) µA mg^−1^ for the electrospun materials. (65.2±7.0) µA mg^−1^ corresponds to about 47 fA per cell (calculated using a dry weight of 716 pg/cell determined with the dry weight of an MR-1 filter cake from an anaerobic culture with known cell density). This value is well in line with values reported by Lu et al. [2017] for MR-1 single-cell current production of 165 fA under aerobic conditions to 36 fA at 0.42 mg L^−1^ dissolved oxygen. In this study, the anode was poised at ∼200 mV (sat. Ag/AgCl, 400 mV vs. NHE), and the cells were quantified by confocal laser scanning microscopy. With real single-cell probing using optical tweezers, a current of 50 fA at the same potential as this work (200 mV vs. NHE) was determined [Liu et al., 2010]. The lower value of GFD 2 could be explained by interstitial cells that populate the pore space [Shroff et al., 2017] and do not contribute to the current production due to lacking electrical contact to the anode. This effect would be less pronounced under micro-aerobic conditions. Interestingly, the current production of single *Geobacter sulfurreducens* cells of (93±33) fA [Jiang et al., 2013] is in the same range as single MR-1 cells.

**Figure 2:**
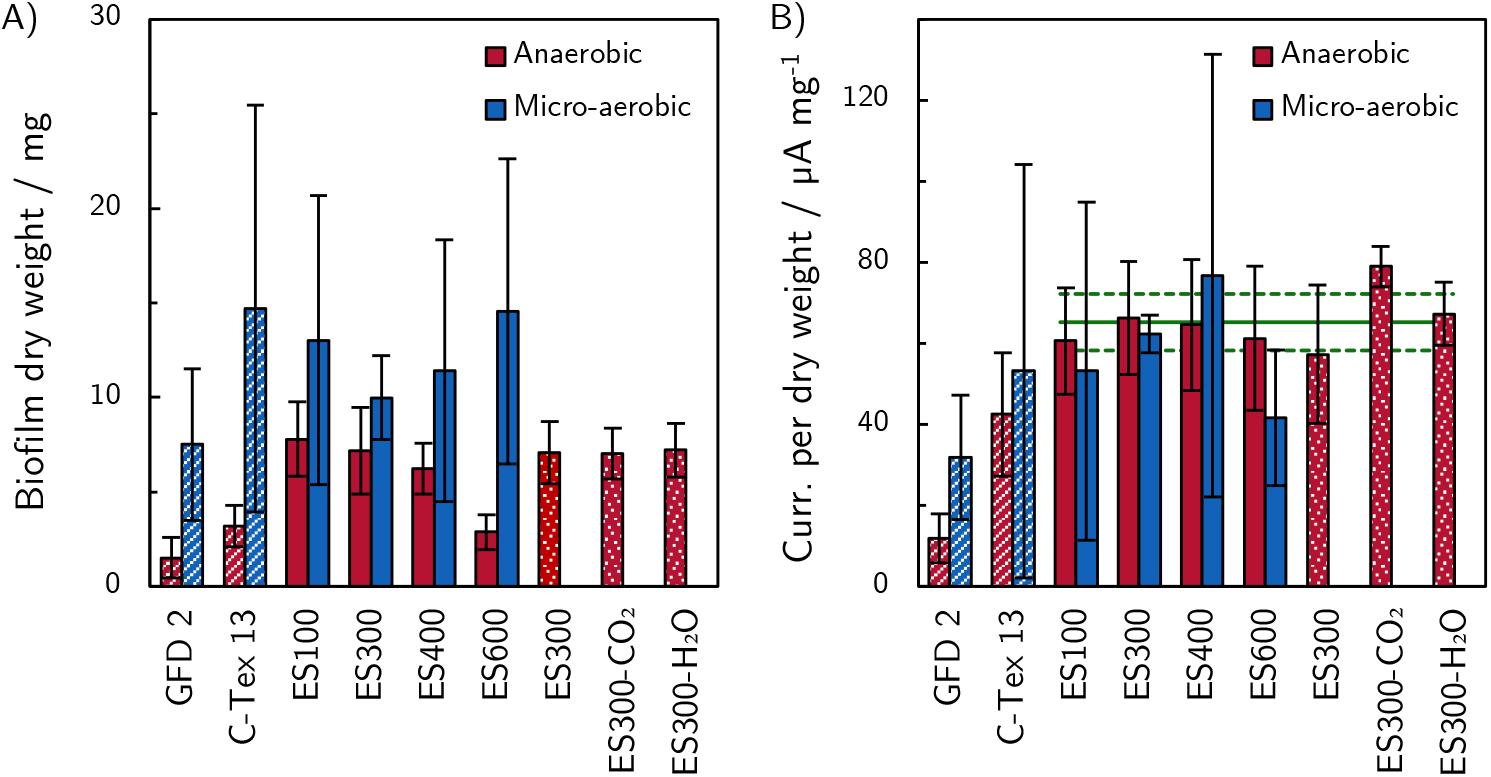
A) Biofilm dry mass found on the anodes after two weeks of operation. B) Final current after two weeks of operation normalized to the biofilm dry weight. The similar values of (65.2± 7.0) µA mg^−1^, as indicated by the horizontal lines in green, found for the electrospun materials under anaerobic and micro-aerobic conditions indicate a proportional relationship between the biofilm found on the electrodes and the current production. The hatched filling indicates the commercial reference materials. All values represent arithmetic means and sample standard deviation and are calculated based on a triplicate.

### 3.5 Morphology of the MR-1 biofilms

SEM images of the biofilm grown under anaerobic conditions on the electrospun anode materials (Fig. 3) reveal biofilm coverage independent of the fiber diameter at the anode front and a trend towards higher biofilm coverage with decreasing fiber diameter at the anode back. The biofilms reaching further into the electrospun anode material and increased bacterial dry weight suggests that thin fiber materials provide a more attractive habitat for MR-1 cells than thick fiber materials. However, under micro-aerobic conditions, a different trend is visible: Higher biofilm coverage on the anode front compared to the anode back is observed. The likely explanation for this observation is oxygen tension towards the anode front due to the abundance of trace amounts of oxygen in the medium [Kim et al., 2016]. It is important to note that the SEM images do not show the biofilm structure in operation. During sample preparation, the drying process leads to a collapse of the porous biofilm onto the anode surface and shrinkage of the cells themselves.

**Figure 3:**
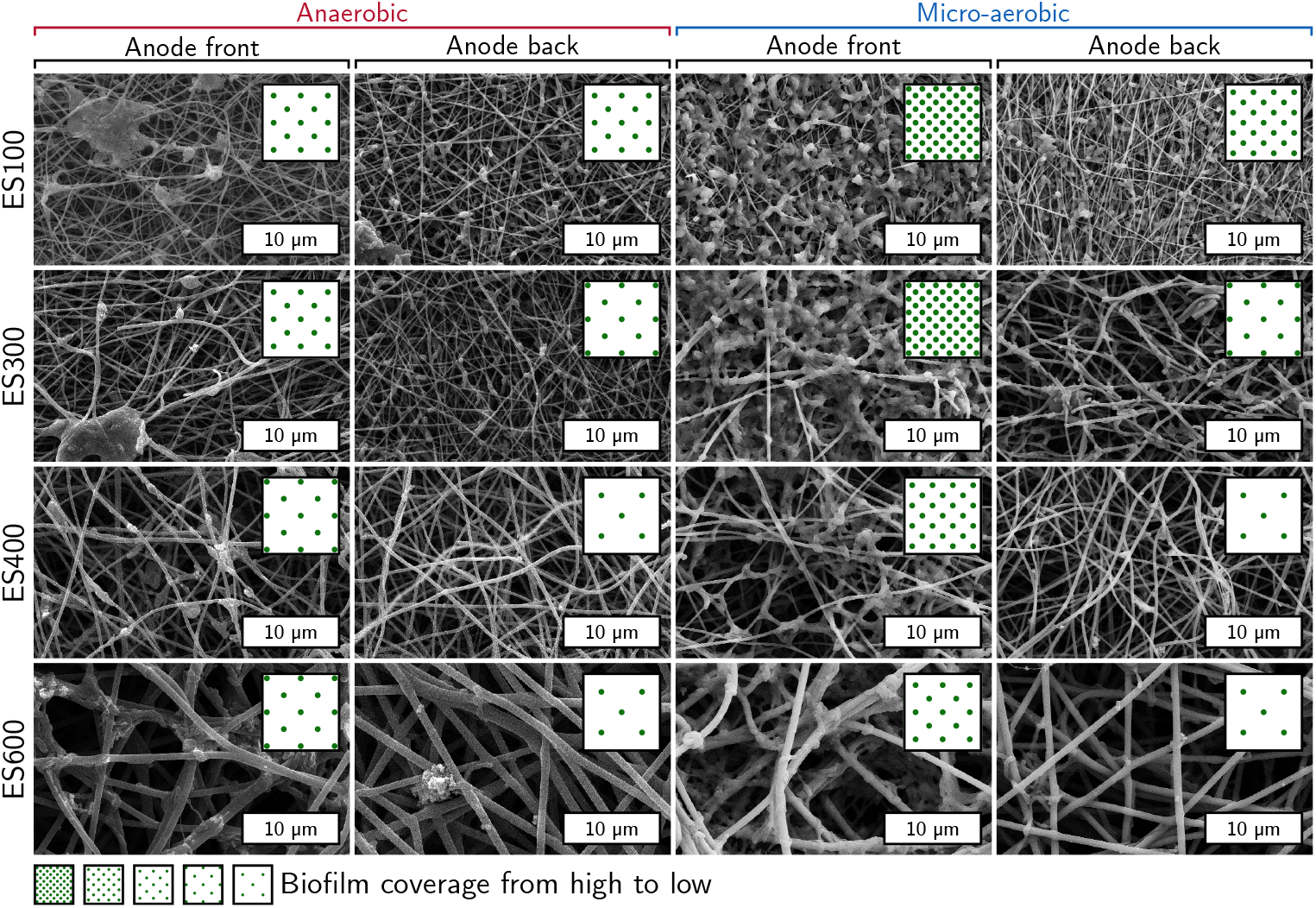
SEM photographs of the biofilms on the electrospun materials and semi-quantitative analysis of the biofilm coverage. Note: the commercial reference materials have been excluded from the biofilm morphology analysis due to major differences in the material morphology. SEM images of the biofilm coverage on these materials are depicted in Fig. S2.6

### 3.6 Total cell growth yield, flavin accumulation, and coulombic efficiency

The planktonic cell growth and flavin accumulation differ significantly under anaerobic and micro-aerobic conditions, as shown in Fig. 5. The presence of trace amounts of oxygen results in cell growth with *OD* _600_ values up to about 0.2. Under anaerobic conditions, the planktonic cell density remains stable, with *OD* _600_ values between 0.03 and 0.05. The total dry mass equivalent of planktonic cells and biofilm on the anodes is depicted in Fig. 4. Only little cell growth with 55 % more bacterial dry mass (biofilm plus the equivalent of planktonic cells) is observed after 14 days of operation. In the case of microaerobic operation, about 380 % more dry mass equivalent accumulates in the reactor after 14 days. A higher fraction, (57 ± 11) %, of the dry weight, is found on the anode under anaerobic conditions compared to (34 ± 17) % under micro-aerobic conditions. The higher cell growth under micro-aerobic conditions is accompanied by flavin accumulation to a final concentration of ∼ 1 µM after 14 days. Under anaerobic conditions, only ∼ 0.1 µM is observed. Considering the similar current to biofilm dry weight ratios of (65.2± 7.0) µA mg^−1^ under micro- and anaerobic conditions (see Section 3.4), it is reasonable to assume that planktonic cells do not contribute considerably to the current production. Hence, mediated electron transfer (MET) by flavins does not significantly affect the overall EET. The low cell growth under anaerobic conditions is surprising because the anode potential in the present work (200 mV vs. NHE) is high enough to drive the DET via OmcA at −110 mV vs. NHE and MtrC at −150 mV vs. NHE as well as MET via free flavins at −250 mV vs. NHE (NHE) [Okamoto et al., 2014]. Higher anode potentials may be preferential for the cell growth energetics and flavin secretion.

**Figure 4:**
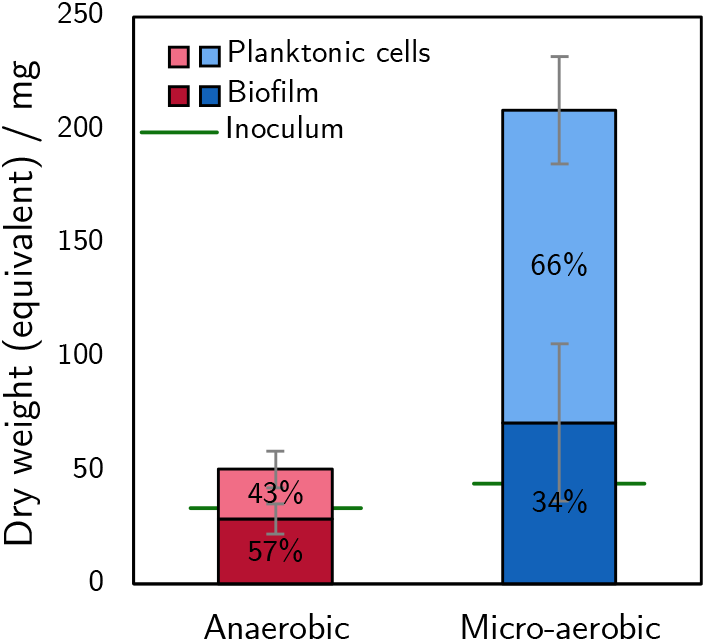
Total cell growth yield under anaerobic and micro-aerobic conditions after 14 days of operation. The growth yield under anaerobic conditions is 55 % and under micro-aerobic conditions 381 %. About 57 % of the total dry mass is found in the biofilm under anaerobic conditions and 34 % under micro-aerobic conditions.

**Figure 5:**
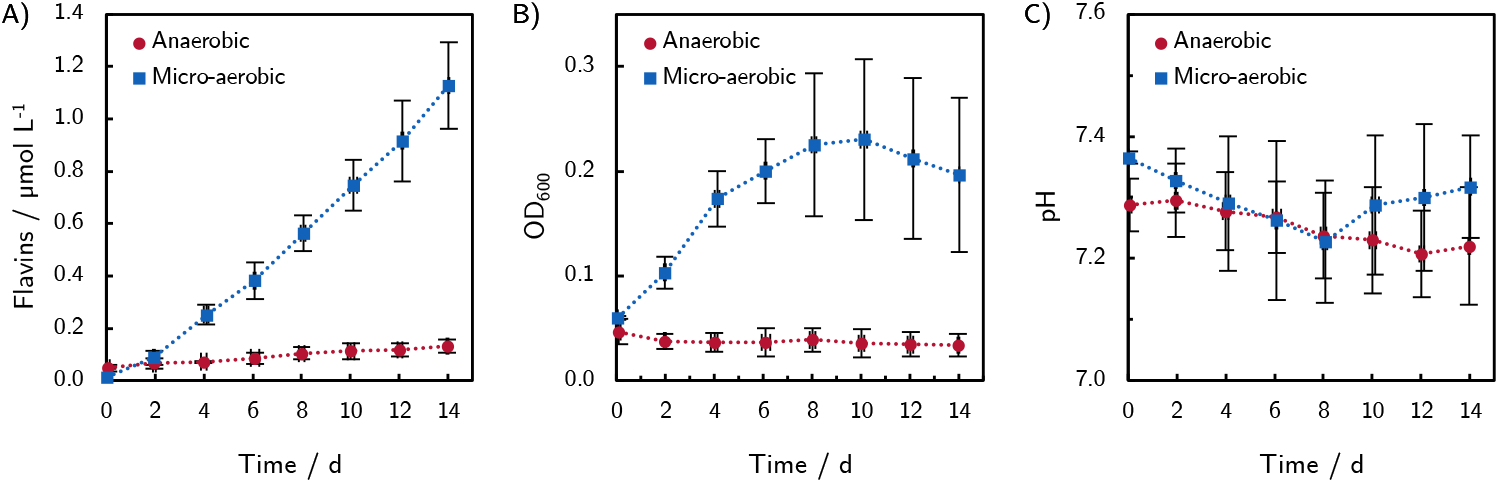
Growth medium characteristics under anaerobic and micro-aerobic conditions. Strongly different behavior can be observed in the flavin accumulation and planktonic cell growth.

For applications such as acetoin production by unbalanced fermentation [Bursac et al., 2017], the coulomb efficiency (CE) is crucial for high product yields. Under anaerobic conditions, the CE reaches values close to unity with (96.6 ± 1.8) %. Under micro-aerobic conditions, the CE drops to (41.6±4.2) %. While the latter value lies in the previously reported range from ∼15 % to ∼75 % [Kipf et al., 2013, Lanthier et al., 2008, Pötschke et al., 2019, TerAvest and Angenent, 2014, Watson and Logan, 2010] for MR-1, the former value exceeds the literature values considerably. Similar high values have only been reported for unbalanced fermentation with engineered *E. coli* and *Shewanella oneidensis* strains [Flynn et al., 2010, Fö rster et al., 2017]. We suspect that our high CE stems from the superior current production with the electrospun anode materials compared to the oxygen leakage into our bioelectrochemical reactor. The pH remained stable under micro- and anaerobic conditions in the range of 7.14 to 7.50 without adjustment.

### 3.7 The role of pore size and surface for the current generation and biofilm formation

As described above, current production is directly linked to the bacterial dry weight attached to the anode. Under anaerobic conditions, the dry weight increases with decreasing fiber diameter of the non-activated electrospun materials. Steam and CO_2_ activation, which introduce micro- and mesopores and increase the surface area, does not further increase biofilm attachment. Therefore, we conclude that the anode materials’ macrostructure determines bacterial attachment to the anode and thus current production. However, as established in Section 3.3 *SRF*, porosity, macropore diameter, and fiber diameter are inseparably linked in fibrous materials on the macroscale. Thus, the higher current production of the electrospun materials with decreasing fiber diameter, which is directly related to the biomass attached to the anode, cannot be attributed directly to either the increased *SRF* or the reduced macropore size. The literature does not provide an in-depth analysis of macropore size and the related surface regarding current production and biofilm formation. In the following, we discuss microscopic processes within biofilms and electrodes that allow attributing the high current production of macroporous electrospun materials to either pore size or surface. Electrospun fiber materials provide high *SRF* s for electron discharge and mean pore diameters in the range of 0.4 µm to 1.6 µm (Tab. 1) allow the bacteria to penetrate the anode pore space and attach to the internal surface. Chong et al. [2019] concluded that pore sizes in the order of a millimeter are preferential for the current production with bacteria that form dense biofilms such as the *Geobacter spp*. and sludge-derived mixed communities. Smaller pores lead to (partial) pore-clogging that hinders the mass transport of nutrients and metabolic products and eventually limits the achievable current densities. MR-1 forms loose biofilms that do not block the pores and do not cover the entire fiber surface (Fig. 3). Hence, the surface area is unlikely to limit bacterial attachment and current production. This raises the question of whether the confinement of cells in proximity to the anode surface or a more attractive chemical microenvironment within the anode pore space leads to increased biofilm formation with decreasing fiber diameter. A recent study by Pirbadian et al. [2020] revealed that the cell membrane hyperpolarization, a measure for metabolic activity, decreases rapidly with the distance from the anode under anaerobic conditions. The membrane hyperpolarization drops within (8.9±12.4) µm to 1*/*e of the value in contact with the electrode. Upon addition of 5 µM riboflavin, this value increases to (21.8±4.3) µm. Biofilms that were grown on flat graphite reach a similar thickness [Kitayama et al., 2017]. Therefore, pores, larger than a few µm, will be occupied by the biofilm with partially reduced average metabolic activity and current production per cell (panel A in Fig. 6). The low current per dry weight values for GFD 2 under anaerobic conditions (little flavin accumulation) and the increase under micro-aerobic conditions (flavin accumulation in the range of 1 µM) can thus be attributed to the large pores of GFD 2. Three mechanisms could explain the decreasing metabolic activity with increasing distance from the anode surface: consecutive redox-cycling of adjacent cells [Okamoto et al., 2012], MET through self-secreted flavins [Carmona-Martinez et al., 2011, Marsili et al., 2008], and a dynamic biofilm structure caused by motile cells that increase the apparent surface coverage with cells, enabling more cells to participate in EET [Okamoto et al., 2012]. Therefore, we cannot attribute the high current densities of electrospun materials to either of the above EET mechanisms. The mean pore diameters of the electrospun materials of 0.4 µm to 1.6 µm are in the same range as the MR-1 cell size (2 µm to 3 µm in length and 0.4 µm to 0.7 µm in diameter, [Venkateswaran et al., 1999]). Therefore, it is not surprising that the current per dry weight for all electrospun materials is similar, considering that the cells must be in close contact with the anode surface in small pores. The average value of 47 fA per cell, we determined in Section 3.4, may represent the metabolic limit of MR-1 cells under this study’s experimental conditions. The total cell volume in the anode is about 8.5 mm^3^ (estimated from the volume of a single cell 0.7 µm^3^), the maximum current density (255 µA cm^−2^, and 47 fA). This is about 8 % of the pore space of the electrospun materials. Therefore, the available pore space is not limiting biofilm formation. A possible explanation for the low cell density within the pore space could be a self-inhibiting effect caused by gradients of metabolic products: i.e., protons and the related local acidification inside the anode (panel B in Fig. 6). Presumably, redox-cycling of flavins occurs between the anode and the MR-1 cells (panel C in Fig. 6). Increasing numbers of MR-1 cells inside the anode shift the equilibrium of oxidized and reduced flavins towards the reduced form. Therefore, the positive chemotaxis of MR-1 cells to oxidized riboflavin [Kim et al., 2016] is decreased by the presence of MR-1 cells. In turn, both smaller pore diameter and larger surface area increase the regeneration of oxidized flavins. Thus, a gradient of oxidized flavins can be sustained with larger numbers of cells within the pore space. Small pore diameter (shorter diffusion distances) and the surface (more reaction sites) would enhance this mechanism. Okamoto et al. [2014] proposed a different role of pores concerning flavins: dead-end pores fully covered by cells could lead to a local increase of flavin concentration in the dead-end pores, enhancing the MET (panel D in Fig. 6). Also, small pores would decrease diffusion distances. This proposed effect requires dead-end pores with sizes within close boundaries: The lower limit is set by the molecular size of flavins (1.3 nm^2^ to 1.5 nm^2^, [Zou et al., 2016]), and the cell footprint gives the upper limit in the order of 1 µm^2^. We cannot exclude this possibility, as the present study’s activated materials mainly feature micropores and open macropores. In conclusion, we can categorize the above processes in surface-related processes (cellular attachment, DET, and oxidation of flavins) and transport processes (flavins, electrons in the biofilm, and bacteria). As MR-1 biofilms do not clog pores, transport processes can be expected to be accelerated by small macropores. Materials with small macropores also provide a larger surface close to the surrounding bulk medium and allow steeper gradients that enhance mass transport of nutrients, oxygen, and metabolic products.

**Figure 6:**
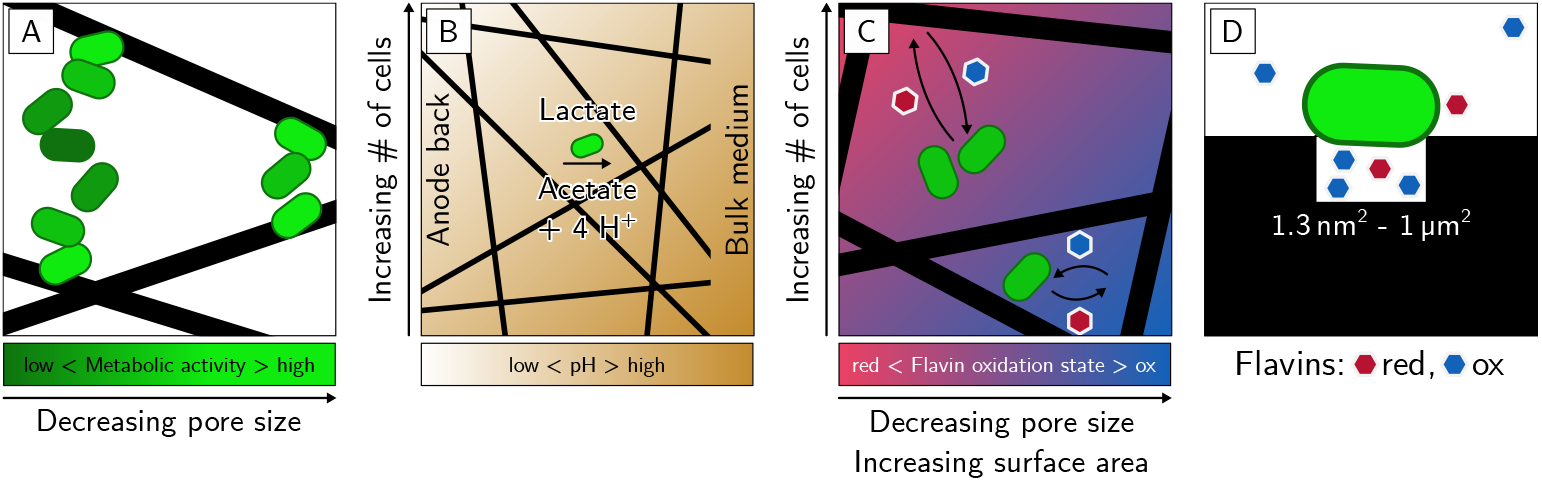
Illustration of microscopic processes in the biofilm and their dependence on pore size, surface area, and the number of cells inside the anode. A) Metabolic activity as a function of the distance to the closest anode surface (free adaption of Pirbadian et al. [2020]. B) Acidification in the anode as a function of distance from the bulk medium and cell density. C) Flavin oxidation state as a function of pore size and cell density inside the anode. D) Flavin accumulation in dead-end pores, according to Okamoto et al. [2014].

### 3.8 Implications for further research

Our material ES300 with open 0.78 µm macropores exhibits a maximum current density of (255 ±71) µA cm^−2^, which corresponds to a 1.4-fold increase compared to the previous literature [Pötschke et al., 2019]. In addition, this extraordinarily high current density is accompanied by a coulombic efficiency (CE) of (96.6±1.8) % that exceeds the previously reported values considerably (see Section 3.6). Both, high CE and current density are required for commercial applications with a high product yield and production rate. As shown by our experimental results, the high current production is linearly related to the bacterial dry weight found on the anodes. The electrode surface is only partially covered with cells, suggesting that the surface area does not directly limit cell attachment. Also, the pore space of the electrospun material is only filled to about 8 % with cells. The underlying mechanisms, that determine the attractiveness of the anodic habitat for MR-1 are not fully understood, and require further research. Further efforts to improve current densities with MR-1 should therefore focus on enhanced biofilm formation. Possible strategies include the chemical modification of the carbon surface, tailoring of pore size and shape. We discussed processes that affect the chemical microenvironment inside the anode material and might cause the improved biofilm formation with decreasing fiber diameter:

- Cellular attachment, DET, and oxidation of flavins are enhanced by large anode surface areas.
- Mediator transport by diffusion and electron transport in the biofilm profit from small macropores.
- High porosity and interconnected pores are required for the transport of nutrient and metabolic products between the bulk medium and the anode’s interior.

Besides the material development, the chemical microenvironment inside the anode could be improved through media optimization, e.g. increase in buffer capacity. Special care needs to be taken during material characterization to ensure anaerobic conditions. Even trace amounts of oxygen lower the CE considerably and (partially) mask the macrostructure’s effect on the current production. Planktonic cell growth and flavin accumulation are indicators for oxygen contamination.

## 4 Conclusions

The current production of Shewanella oneidensis MR-1 with tailored electrospun anode materials with fiber diameters between 108 nm to 623 nm was characterized. Our highest current production of (255 ± 71) µA cm^−2^ with the 286 nm fiber diameter material exceeds the highest value reported in the literature 1.4-fold. The high current density is accompanied by a coulombic efficiency of (96.6±1.8) %. Additional activation did not improve the current production significantly. We observe a linear relationship between current production and bacterial dry weight and increased penetration depth of the biofilm. We conclude that the anode material’s attractiveness for MR-1 cells is most likely determined by both the macropore size and the accessible surface area.

## 5 Experimental Section

### 5.1 Preparation of tailored electrospun and commercial reference anodes

Four electrospun carbon fiber mat materials with average fiber diameters of 108 nm (ES100), 286 nm (ES300), 400 nm (ES400), and 623 nm (ES600) were fabricated by electrospinning of PAN with concentrations between 6 wt% and 13 wt% in N,N-dimethylacetamide and subsequent carbonization at 1100 °C as previously reported [Erben et al., 2021]. Activation was conducted for 4 h with CO_2_ at 832 °C and with steam at 757 °C. Two commercial carbon materials, knitted steam activated carbon C-Tex 13 (Mast Carbon International Ltd., United Kingdom) and graphite felt GFD 2 (SGL Carbon SE, Germany), were used as reference materials [Dolch et al., 2014, Kipf et al., 2013, 2014] for comparison. SEM photographs of these materials are provided in Fig. S1.1. The anodes were cut into pieces of 2 cm^2^× 2 cm^2^ and mounted in polycarbonate frames exposing 2.25 cm^2^ to the growth medium (Fig. S1.2). To remove gas bubbles from the porous anode structures, the anode material was wetted with isopropyl alcohol, rinsed with DI water, and placed in vacuum for approximately 10 minutes before the experiments.

### 5.2 Anode material characterization

The electrochemically active surface (*ECAS*) of the anodes was determined via the measurement of the double-layer capacitance using a Solartron Analytical potentiostat (1470E, Farnborough, United Kingdom) as reported previously [Erben et al., 2021]. Nitrogen adsorption was conducted at 77 K using a Belsorp-Max instrument (BEL Japan Inc., Japan) for the electrospun materials and a Sorptomatic 1990 (CE Instruments, Italy) for the commercial reference materials. Scanning electron microscopy (SEM) was performed using a DSM 962 (Carl Zeiss Microscopy GmbH, Germany) and an SX 100 (Cameca, France). The macropore sizes were estimated from SEM photographs using the software ImageJ and DiameterJ [Hotaling et al., 2015] as described in Section S4 in the supplementary information. The porosity *φ* was calculated based on the anode thickness in the mounting frames *t* according to *φ* = 1 − *ρ*_A_ (*tρ*)^−1^. Herein, *ρ*_A_ is the material’s area density, and *ρ* is the density of PAN-derived carbon (1.75 g cm^−3^, [Wangxi et al., 2003]). The internal surface area was assessed by the *ECAS* method [Erben et al., 2021]. From the weight-specific *ECAS* and *S* _BET_ surface area, and the area density *ρ*_A_, the Surface Roughness Factor (*SRF*) was calculated according to *SRF*_ECAS_ = *ECAS* · *ρ*_A_ and *SRF*_BET_ = *S*_BET_ *ρ*_A_. The *SRF*, defined by the ratio of the internal surface area and the anode’s projected surface area, allows the direct comparison of the materials with different area densities. Similarly, the absolute micro-and mesopore volume *V* _Micro_ and *V* _Meso_, determined from N_2_ adsorption isotherms (*t*-plot and BJH analysis respectively), was calculated from the weight-specific value *𝒱* for the projected anode area: *V* = *𝒱* · *ρ*_A_·2.25 cm^2^. The macropores of the electrospun materials and GFD 2 were assessed by digital image processing and 3D modeling as described in Section S4 in the supplementary information. With this method, a 2D SEM image is analyzed for its apparent porosity and pore diameter. In a second step, the 2D apparent pore diameter is converted into a 3D pore diameter by comparison with a 3D model of the fiber material morphology. This unique method was chosen because mercury intrusion porosimetry underestimates the pore sizes of compressible materials Giesche [2006], Rutledge et al. [2009].

### 5.3 Bioelectrochemical characterization

MR-1 cells were prepared following the procedure described by Kipf et al. [2013]. In short, MR-1 cells were first cultivated aerobically in Lysogeny broth (LB medium) and then transferred to a growth medium containing 50 mM Na-D/L-lactate and 100 mM fumarate. The cells were washed with washing buffer (growth medium lacking lactate and fumarate) thrice before inoculation of the bioelectrochemical reactor (Fig. S1.3). The amount of inoculum was adjusted to an initial optical density (*OD* _600_) of ∼0.05. Before inoculation, the bioelectrochemical reactor was sterilized by steam autoclavation at 121 °C for 20 min. The bioelectrochemical reactor contained 1 L anode medium with 50 mM initial concentration of Na-D/L-lactate as an electron donor and no electron acceptor other than 6 anodes. The anodes were polarized at −41 mV against a sat. Calomel reference (SCE, KE11, Sensortechnik Meinsberg, Germany) using PGU-MOD 500 mA potentiostats (IPS Elektroniklabor GmbH & Co. KG, Münster, Germany). To minimize non-coulombic currents, the anodes were polarized at least 12 h before the inoculation. A platinum mesh acted as a counter electrode. The current production was recorded for 14 days. In the present publication, potentials are additionally provided vs. Normal Hydrogen Electrode (NHE) at 25 °C for better comparison. This way, different temperature dependencies of the redox couples are accounted for. The reactor is operated at 30 °C and its headspace is continuously purged with humidified N_2_ at 125 mL min^−1^ for anaerobic conditions (*<*20 ppm). Trace amounts of O_2_ (200 ppm - 400 ppm corresponding to 1.4 µg L^−1^ - 2.8 µg L^−1^ dissolved oxygen) were introduced into the reactor by guiding the gas flow through an O_2_ permeable silicone tube. Oxygen concentrations were determined with a Fibox 3 LCD trace oxygen meter (PreSens Precision Sensing GmbH, Germany). The medium was stirred at 300 rpm and sampled every other day to monitor pH, cell density, lactate consumption, and flavin concentration.

### 5.4 Medium monitoring

Flavin concentration in the medium was measured with a Nanodrop 3300 spectrometer (emission at 525 nm, excitation 470 nm) using riboflavin (RF) as a standard. RF exhibits similar fluorescence signals as FMN and FAD [Valle et al., 2012]. Therefore, the flavin concentrations reported comprise FMN, FAD, and RF. The CE was calculated based on the lactate consumption under microaerobic conditions and based on the acetate accumulation under anaerobic conditions assuming partial oxidation of lactate to acetate with 4 electrons per molecule [Lanthier et al., 2008]. The initial and final lactate and acetate concentrations were determined by high-performance liquid chromatography (Elite La Chrome, Hitachi, Japan) equipped with an Aminex HPX-87H column (Bio-Rad, Germany) and RI-detector (RefraktoMax 521, IDEX Health & Science, Japan). The planktonic dry weight m in the medium was determined by {*m*}_mg_ = 716 · {*V*}_Liter_ · {*OD* _600_}. The conversion factor was determined experimentally with the dry weight of a cell filter cake from medium with known volume and *OD* _600_.

### 5.5 Biofilm analysis

The biofilm dry weight was quantified after 14 days of operation as follows. The anodes were placed in 1 mL of lysis buffer (20 mM Tris/HCl, 16.5 mM Triton X-100, 137 mM NaCl, protease inhibitor cocktail for bacteria 25 µL mL^−1^, Carl Roth, Germany) at 4 °C overnight. The lysate’s protein content was then analyzed using a biuret test according to the instructions (Roti-quant universal, Carl Roth, Germany). Dry mass was calculated by comparison to cell lysate with known dry weight. For SEM imaging, the biofilm was fixated with 2.5 % glutaraldehyde in anode medium lacking lactate for 24 h at 4 °C. The samples were then dehydrated with a series of 10 %, 30 %, 50 %, 70 %, 80 %, 90 %, and 100 % ethanol in washing buffer for 10 min each and air-dried at room temperature.

## 6 Acknowledgements

We are grateful for the financial support from the German Ministry of Education and Research (BMBF) under the program 03SF0496A. We thank Dr. Guillaume Alexis Adrien Pillot for fruitful discussions and critical feedback during the writing process.

