## Supplemental Material for "High current production of *Shewanella oneidensis* with electrospun carbon nanofiber anodes is directly linked to biofilm formation"

February 5, 2021

The related publication can be accessed via `http://doi.org/updateDOI/`.

Preprint submitted to ChemElectroChem

#### Contents

**S1 Anode materials and bioelectrochemical reactor** **S2**

**S2 Supplemental figures** **S4**

**S3 Analysis of the adsorption and desorption isotherms** **S6**
S3.1 Brunauer-Emmet-Teller (BET) method . . . S6
S3.2 Barret-Joyner-Halenda (BJH) method . . . S6
S3.3 Harkins-Jura *t*-plot . . . S6
S3.4 Material properties (extended) . . . S10

**S4 Pore diameter estimation** **S11**
S4.1 Geometrical model (2D) . . . S11
S4.1.1 Macropore diameter estimation . . . S11
S4.1.2 Surface roughness factor estimation . . . S12
S4.1.3 Correlation of surface roughness factor *SRF* and fiber diameter $d_F$ . . . S12
S4.2 Macropore diameter estimation using a 3D model . . . S13
S4.2.1 SEM image analysis . . . S13
S4.2.2 3D macropore model . . . S14
S4.2.3 Model construction . . . S14
S4.2.4 Apparent pore diameter and porosity . . . S15
S4.2.5 3D continuous pore diameter distribution . . . S16
S4.2.6 Conversion of apparent pore diameter to 3D pore diameter . . . S16

**S5 Literature analysis** **S17**

**References** **S20**

#### S1 Anode materials and bioelectrochemical reactor

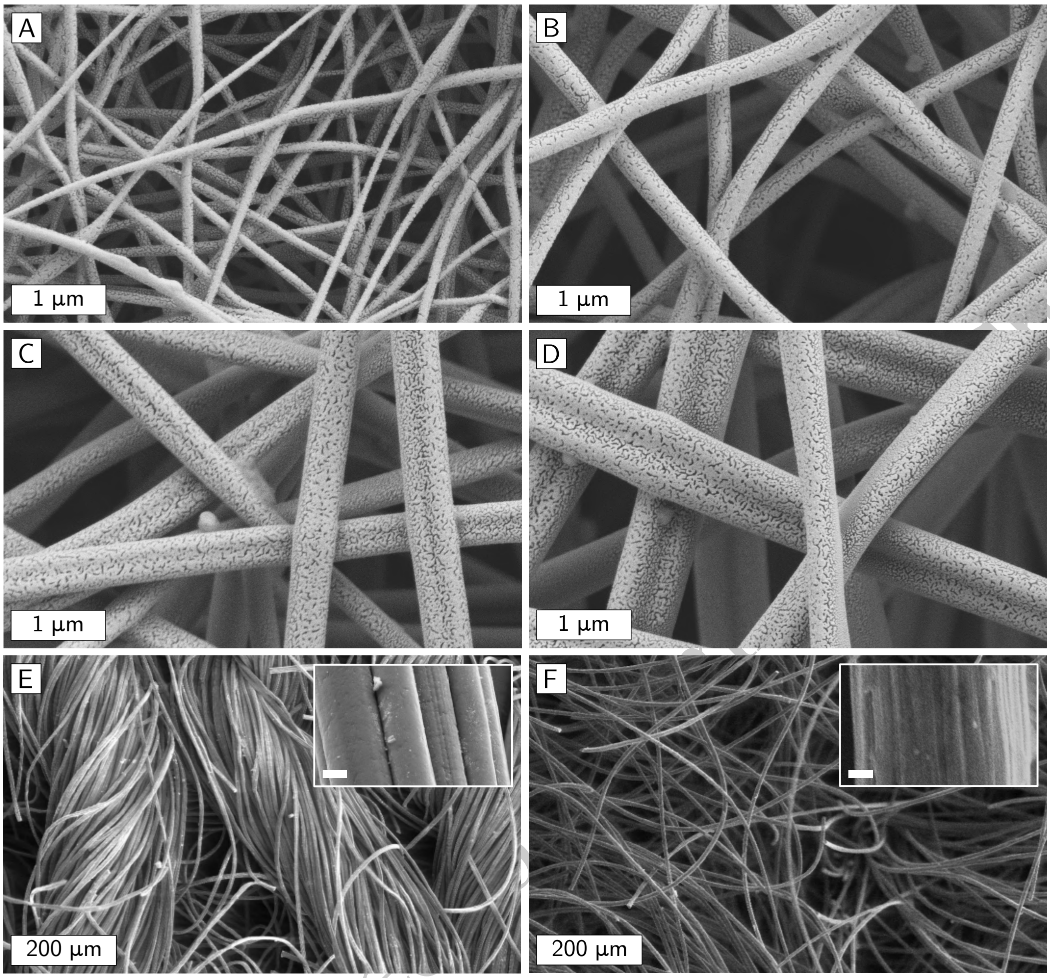


**Fig. S1.1:** SEM images of the anode materials. A-D: Electrospun carbon fiber materials ES100, ES300, ES400, and ES600. E: Knitted steam activated carbon C-Tex 13 (Mast Carbon International Ltd., Hampshire, United Kingdom). F: Graphite felt GFD 2 (SGL Carbon SE, Wiesbaden, Germany). The insets show a magnification of the individual fibers (scale bar 1 µm).

Preprint submitted [illegible]

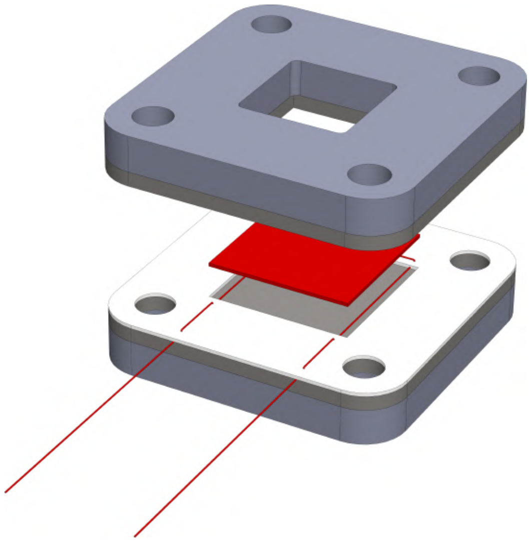

**Fig. S1.2:** Rendering of the electrode assembly. The electrode material 2x2 $cm^2$ in size and the titanium wire are depicted in red. The electrode material is sandwiched between silicone sheets together with a 0.5 mm thick spacer with a $2\,cm^2 \times 2\,cm^2$ cutout. The area exposed to the medium is $1.5\,cm^2 \times 1.5\,cm^2$.

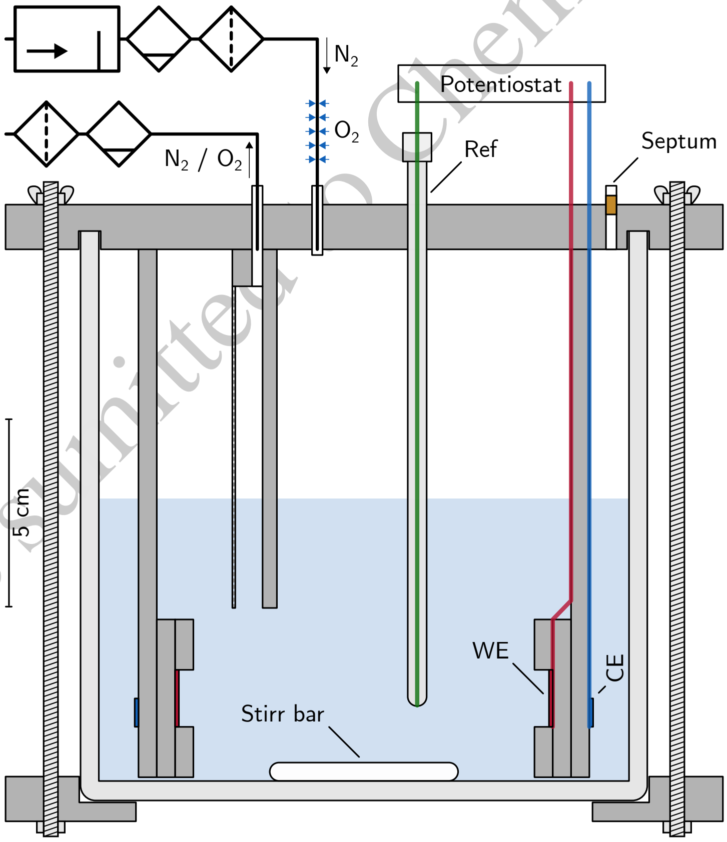


**Fig. S1.3:** Schematic of the bioelectrochemical reactor with electrical peripherals and gas supply system. Note: The optional oxygen permeable silicone tubing is depicted as blue arrows pointing towards the $N_2$ supply. The working electrode is connected to the potentiostat in a two-wire configuration in order to reduce the uncompensated resistance. Abbreviations: working electrode (WE), counter electrode (CE), reference electrode (Ref).

Preprint submitted to ChemElectroChem

#### S2 Supplemental figures

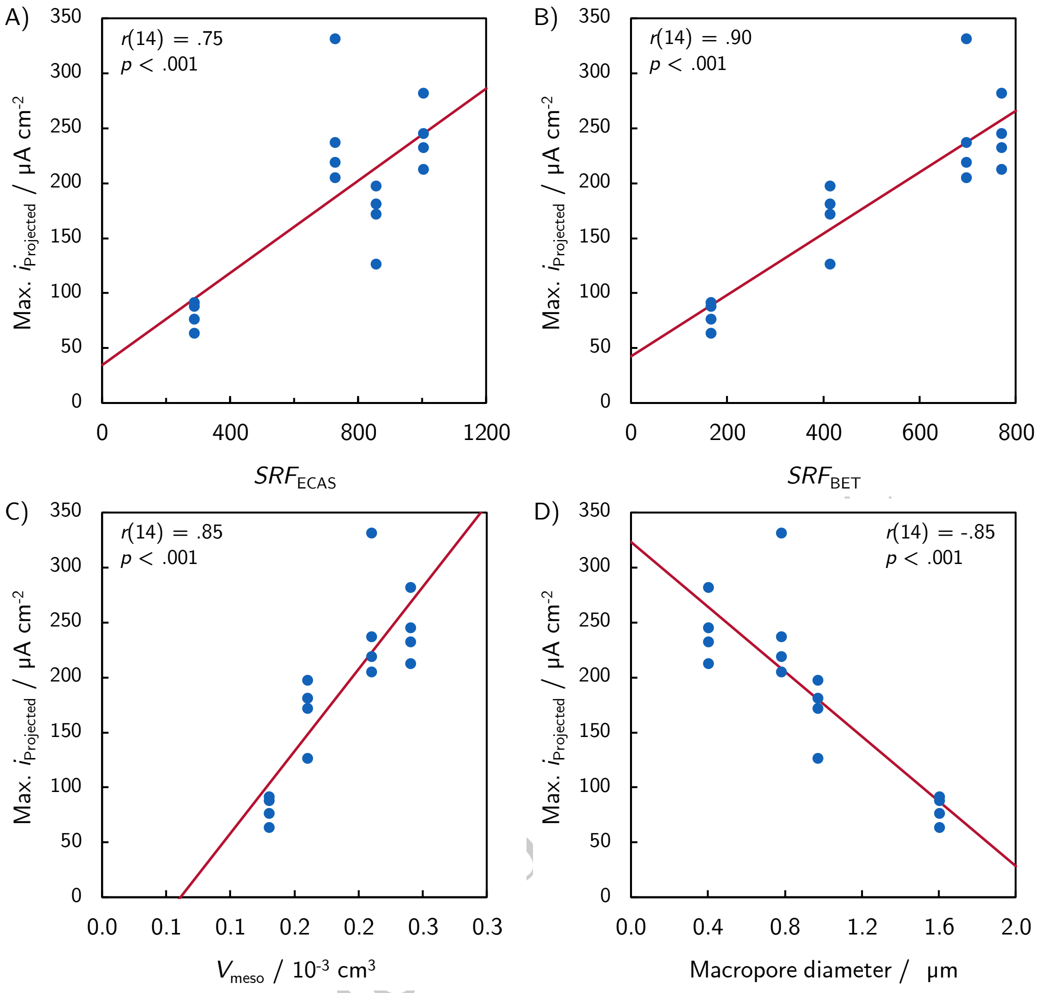


**Fig. S2.4:** Linear correlation plots of the maximum projected current density with the material properties surface roughness factor ($SRF_{ECAS}$ and $SRF_{BET}$), mesopore volume $V_{meso}$, and macropore diameter.

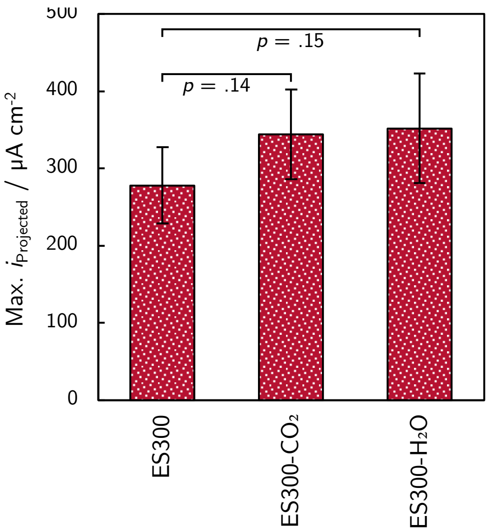


**Fig. S2.5:** Comparison of the maximum projected current density of ES300, ES300-$CO_2$, and ES300-$H_2O$. The $p$-values were calculated with a Welch corrected one tail $t$-test.

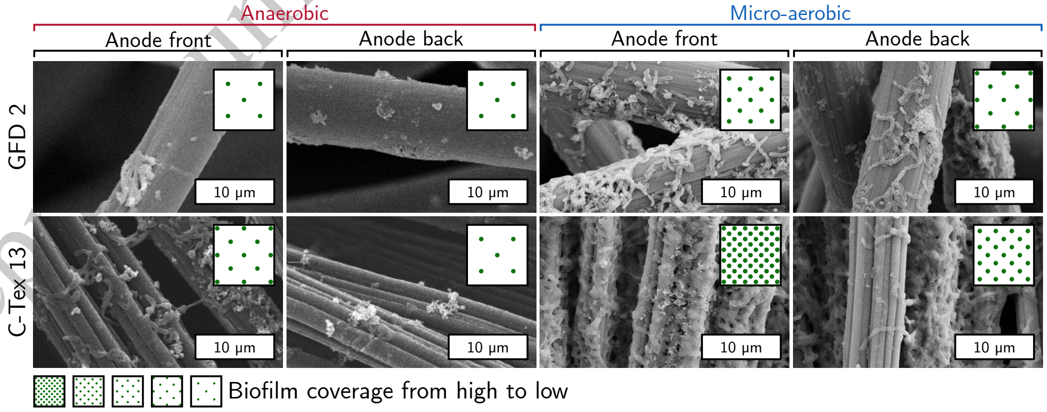


**Fig. S2.6:** SEM images of the biofilms on the commercial reference materials.

Preprint submitted to ChemElectroChem

#### S3 Analysis of the adsorption and desorption isotherms

All materials were characterized by nitrogen adsorption. The specific BET surface ($SSA_{BET}$), the meso- and micropore volume given in Tab. S3.1 were calculated from the adsorption- and desorption isotherms as described below.

##### S3.1 Brunauer-Emmet-Teller (BET) method

The BET method [1] yields a measure for the surface area by linearization of nitrogen adsorption isotherms:

$$\frac{1}{V_{\mathrm{ads.}}} = \frac{c-1}{V_{\mathrm{m}}c}\left(\frac{p}{p_0}\right) + \frac{1}{V_{\mathrm{m}}c}. \tag{S3.1}$$

Here, $V_{\mathrm{ads.}}$ is the adsorbed quantity of nitrogen and $c$ is the BET constant. $V_{\mathrm{ads.}}$ can be calculated from the $y$-axis intercept $d$ and the slope $k$:

$$V_{\mathrm{m}} = \frac{1}{k+d}. \tag{S3.2}$$

With the Avogadro constant $N_{\mathrm{A}}$, the adsorption cross section $s$, and the molar volume $v_{\mathrm{m}}$ of nitrogen the specific surface area is given by:

$$S_{\mathrm{BET}} = \frac{V_{\mathrm{m}} N_{\mathrm{A}} s}{v_{\mathrm{m}}}. \tag{S3.3}$$

The BET linearizations are depicted in Figs. S3.7, S3.7, and S3.9 in panel B.

##### S3.2 Barret-Joyner-Halenda (BJH) method

The BJH mesopore size distribution [2] is obtained by iterative analysis of the desorption branch under the assumption of cylindrical pores and that the adsorbent is retained in the pores either by physical adsorption on the walls of the pores or by capillary condensation. The poresize distributions calculated using the Belsorp Adsorption/Desorption Data Analysis Software and Sorptomatic Advanced Data Processing (commercial materials) are given in Figs. S3.7, S3.7, and S3.9 in panel C.

##### S3.3 Harkins-Jura *t*-plot

According to Harkins and Jura [3], the relative pressure $p/p_0$ can be related to a statistical thickness of the fluid layer $t_{\mathrm{Harkins-Jura}}$ in nm:

$$t_{\mathrm{Harkins-Jura}} = 10 \cdot \sqrt{\frac{13.99}{0.34 - \log(p/p_0)}}. \tag{S3.4}$$

The micropore volume is given by the volume of adsorbed nitrogen at zero thickness of the layer calculated by linear regression of the *t*-plot (Figs. S3.7, S3.7, and S3.9 in the panel D).

Preprint sumitted to ChemElectroChem

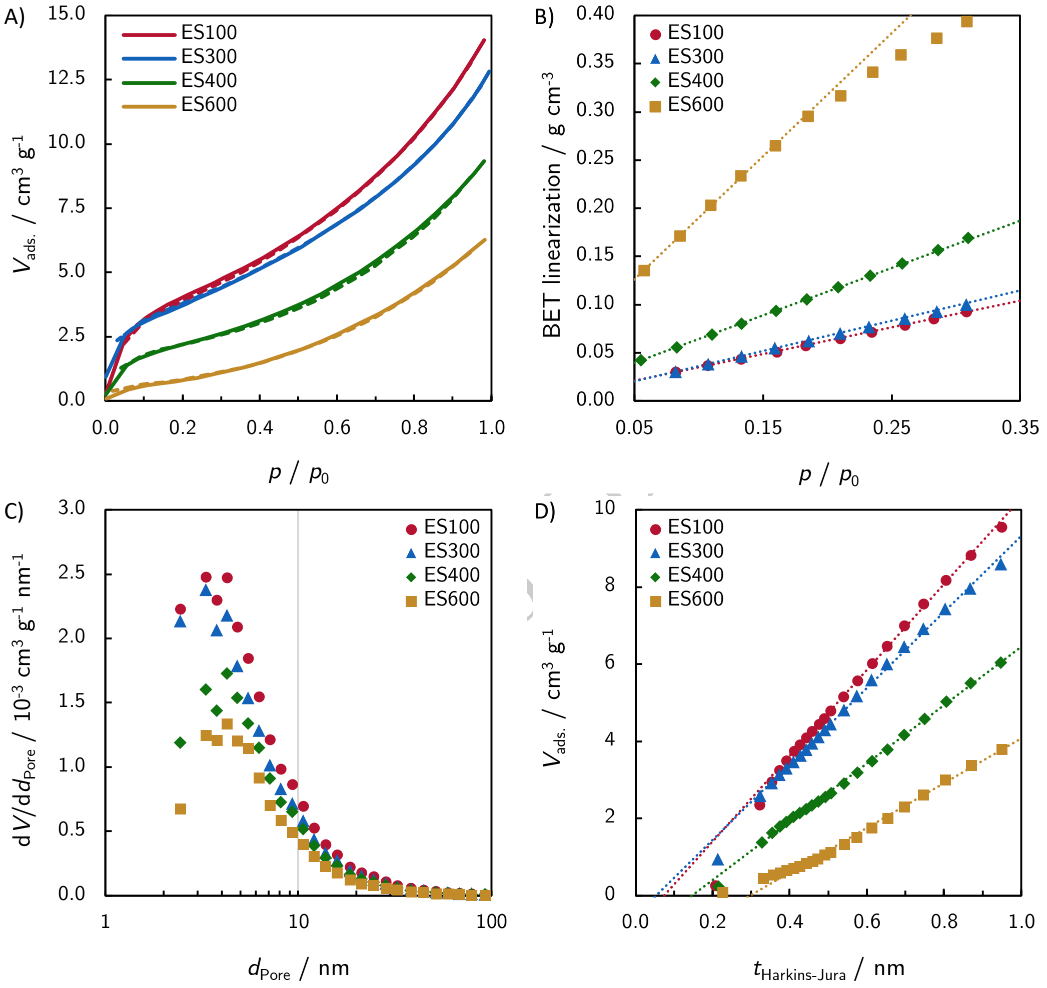


**Fig. S3.7:** Nitrogen adsorption measurements of the non-activated electrospun materials. A) adsorption (solid lines) and desorption (dashed lines) isotherms, B) BET linearization, C) Harkins-Jura *t*-plot for micropore volume determination, D) BJH mesopore size distribution. The dotted lines represent linear regressions used for the evaluation of the BET surface and the micropore volume.

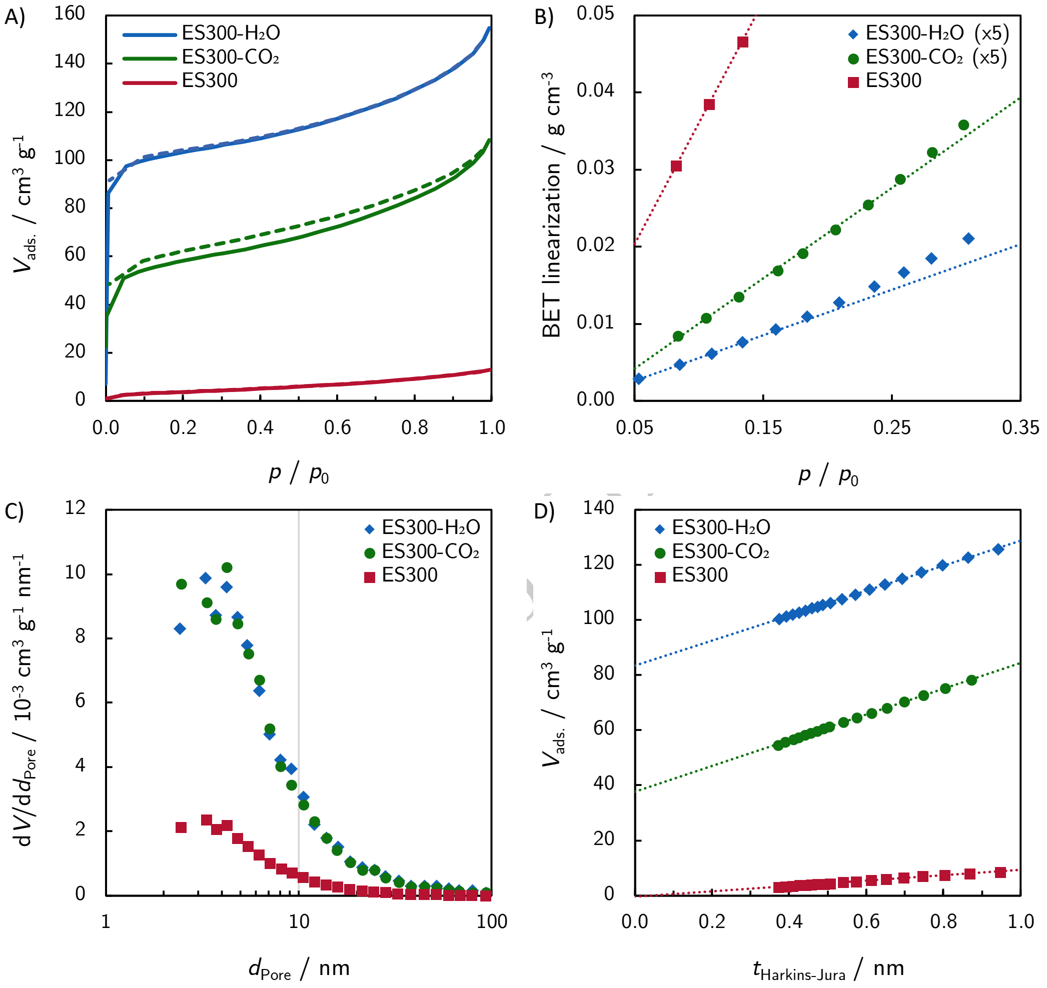


**Fig. S3.8:** Nitrogen adsorption measurements of the non-activated and activated ES300 materials. A) adsorption (solid lines) and desorption (dashed lines) isotherms, B) BET linearization, C) Harkins-Jura *t*-plot for micropore volume determination, D) BJH mesopore size distribution. The dotted lines represent linear regressions used for the evaluation of the BET surface and the micropore volume.

Preprint ... chem

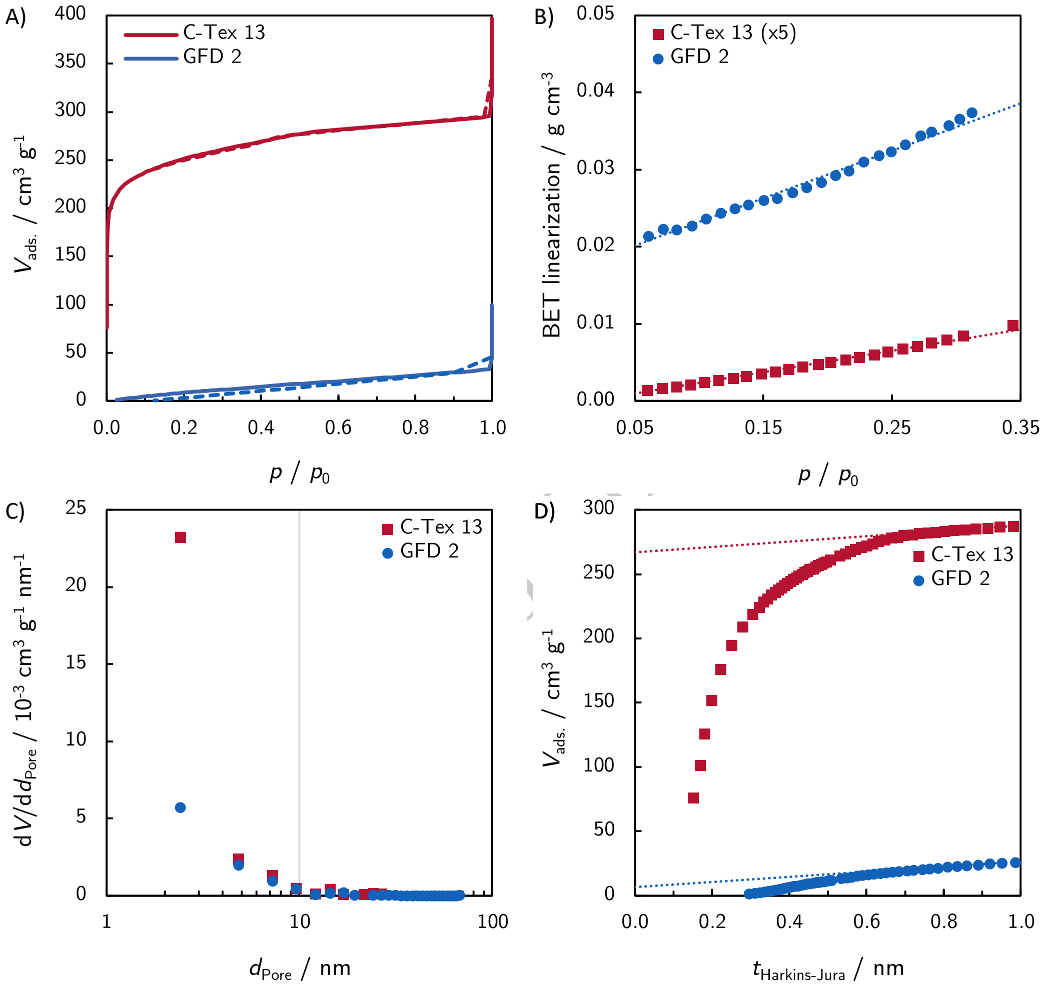


**Fig. S3.9:** Nitrogen adsorption measurements of the commercial reference materials C-Tex 13 and GFD 2. A) adsorption (solid lines) and desorption (dashed lines) isotherms, B) BET linearization, C) Harkins-Jura *t*-plot for micropore volume determination, D) BJH mesopore size distribution. The dotted lines represent linear regressions used for the evaluation of the BET surface and the micropore volume.

Preprint ... ChemRxiv

Preprint sumitted to ChemElectroChem

#### S3.4 Material properties (extended)

**Table S3.1:** Extended properties of the investigated anode materials.

| Material | Diameter / nm | Area density / g m$^{-2}$ | $v_{Micro}$ / 10$^{-3}$ cm$^3$ g$^{-1}$ | $v_{Meso}$ / 10$^{-3}$ cm$^{-3}$ g$^{-1}$ | $d_{Macro}$ / µm | $S_{BET}$ / m$^2$ g$^{-1}$ | *ECAS* / m$^2$ g$^{-1}$ | $\varphi$ / % |
|---|---|---|---|---|---|---|---|---|
| GFD 2 | ∼8000 | 182 | 10 | 48 | 31 | 55 | 0.15 | 94 * |
| C-Tex 13 | ∼7000 | 165 | 413 | 91 | – | 800 | 800 | 81 |
| ES100 | 108 | 50 | – | 21.0 | 0.40 | 15.4 | 20.1 | 94 |
| ES300 | 286 | 51 | – | 18.7 | 0.78 | 13.7 | 14.2 | 94 |
| ES400 | 400 | 48 | – | 14.5 | 0.97 | 8.6 | 17.8 | 95 |
| ES600 | 623 | 51 | – | 11.0 | 1.60 | 3.3 | 5.6 | 94 |
| ES300-$CO_2$ | ∼286 | 44 | 58 | 99 | 0.78 | 189 | 427 | 95 |
| ES300-$H_2O$ | ∼286 | 44 | 129 | 100 | 0.78 | 372 | 785 | 95 |

* Value according to manufacturer datasheet.

**Table S3.2:** Summary of the intercorrelations of the material properties $SRF_{ECAS}$, $SRF_{BET}$, macropore diameter, and $V_{meso}$. The corresponding $p$-values given in brackets were calculated with two degrees of freedom.

| | $SRF_{ECAS}$ | $SRF_{BET}$ | Macropore diameter |
|---|---|---|---|
| $SRF_{BET}$ | .82 ($p$ = .18) | – | – |
| Macropore diameter | -.94 ($p$ = .062) | -.96 ($p$ = .039) | – |
| $V_{Meso}$ | .79 ($p$ = .21) | .98 ($p$ = .017) | -.95 ($p$ = .046) |

### S4 Pore diameter estimation

#### S4.1 Geometrical model (2D)

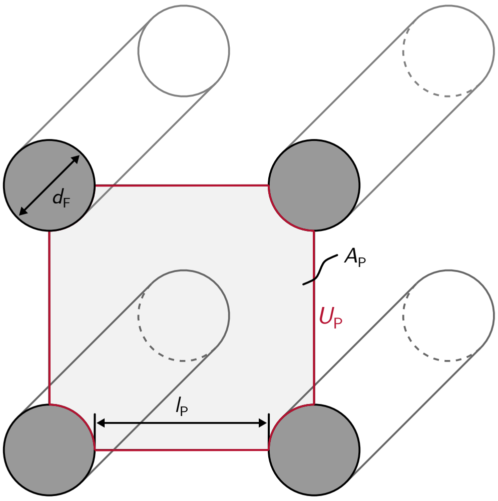


**Fig. S4.1:** Illustration of the fiber material macropore model adapted from Chong et al. [4]. The model is based on an hypothetical cross section through the fiber material.

The macropore model introduced by Chong et al. [4] can be extended for the estimation of the macropore size $d_{\text{Macro}}$ and the surface roughness factor *SRF*. The geometrical model neglects the non-quadratic pore shape of electropun fiber materials and the non-circular cross section of the fibers. The model fails to predict the pore diameter and *SRF* quantitatively but gives a theoretical explanation for the intercorrelations of *SRF*, macropore diameter, and fiber diameter determined experimentally (Tab. S3.2).

##### S4.1.1 Macropore diameter estimation

With the definitions in Fig. S4.1 and the definition for the macropore diameter calculation $d_{\text{Macro}} = 4\,A_P/U_P$ used to evaluate SEM images, one yields an expression for the macropore diameter:

$$d_{\text{Macro}} = \frac{4A_P}{U_P} \approx d_F + l_P.$$

The approximation is valid for highly porous materials with $d_F \ll l_P$. The porosity $\varphi$ can be expressed as

$$\varphi \approx 1 - \frac{\pi d_F^2}{4} \frac{1}{d_{\text{Macro}}^2},$$

and rearranged to the solid fraction in a fibrous material:

$$1 - \varphi \approx \frac{\pi d_F^2}{4 d_{\text{Macro}}^2}. \tag{S4.1}$$

The fiber surface $S_F$ per electrode volume $V_E$ can be expressed as:

$$\frac{S_F}{V_E} \approx \frac{4}{d_F} \frac{\pi d_F^2}{4 d_{\text{Macro}}^2}. \tag{S4.2}$$

Substitution with S4.1 yields

$$\frac{S_F}{V_E} \approx \frac{4}{d_F}(1 - \varphi). \tag{S4.3}$$

Combining S4.2 and S4.3 yields an approximation relating the fiber diameter $d_F$, the pore diameter $d_{\text{Macro}}$ and the porosity $\varphi$:

$$\boxed{d_{\text{Macro}} \approx \underbrace{\sqrt{\frac{\pi}{4(1-\varphi)}}}_{\text{Conversion factor}} \times d_F}\,. \tag{S4.4}$$

Preprint sumitted to ChemElectroChem

The geometrical model overpredicts the pore diameters due to the non-quadratic shape of the pores in electropun fiber materials. However, Eq. S4.4 shows the linear relationship between macropore diameter and fiber diameter found for electrospun fiber materials [5]. The conversion factor will be determined in Section S4.2 with a 3D model.

##### S4.1.2 Surface roughness factor estimation

With the definition of the surface roughness factor *SRF*

$$SRF = \frac{S_\mathrm{F}}{A_\mathrm{E}} = \frac{S_\mathrm{F}}{V_\mathrm{E}} t_\mathrm{E},$$

the projected electrode area $A_\mathrm{E}$, electrode thickness $t_\mathrm{E}$, and the approximation S4.3 one yields a relation between the *SRF* and the fiber diameter

$$\boxed{SRF \approx 4t_\mathrm{E}(1-\varphi) \times \frac{1}{d_\mathrm{F}}}\,. \tag{S4.5}$$

This relation is depicted in Fig. S4.2 together with the *SRF*s determined experimentally by the ECAS and BET methods. While beeing able to predict the general trend towards higher *SRF* with decreasing fiber diameter, the model prediction deviates substantially from the measurements. The deviations of the predicted *SRF* may be explained by the non-circular cross section of the real fibers and sub-diameter porosity that enlarge the fiber surface.

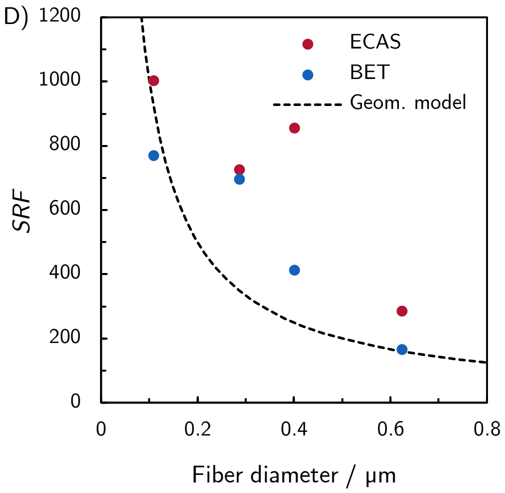


**Fig. S4.2:** Predictions of the geometrical model with 95 % porosity $\varphi$ and an electrode thickness $t_\mathrm{E}$ of 0.5 mm for the *SRF* compared to the experimentally determined values (*ECAS* and BET).

##### S4.1.3 Correlation of surface roughness factor *SRF* and fiber diameter $d_\mathrm{F}$

Combining Eqs. S4.5 and S4.4 yields a correlation for $d_\mathrm{Macro}$ and *SRF* on the macroscale:

$$\boxed{d_\mathrm{Macro} \sim \frac{\sqrt{1-\varphi}}{SRF}}\,. \tag{S4.6}$$

Therefore, it is not possible to modify the macropore diameter without changing the *SRF* for a given electrode thickness and porosity.

Preprint ChemElectroChem

#### S4.2 Macropore diameter estimation using a 3D model

Two-dimensional SEM images do not include depth information required for pore diameter analysis. The following approach allows the conversion of two-dimensional apparent pore diameters to three-dimensional pore diameters by means of a 3D model.

##### S4.2.1 SEM image analysis

To ensure comparable results, the black value threshold of the SEM images used for pore diameter analysis was adjusted manually to limit the depth of focus to about 6 to 7 fiber layers. The apparent pore diameters were then analyzed with *ImageJ* in 4 steps:

1. SEM images were segmented using the *DiameterJ Segment* function [6]. *DiameterJ* performs 24 different segmentation algorithms. An image, that represents best the original was selected for further analysis. A comparison of a original image and the selected segmentation is provided in Fig. S4.1 A) and B).

2. The apparent porosity $\varphi_{\text{Apparent}}$ was calculated by counting white pixels:

$$\varphi_{\text{Apparent}} = \frac{\#\text{ of white pixels}}{\text{Total }\#\text{ of pixels}} \tag{S4.1}$$

3. The apparent pore areas $A_{\text{P}}$ and the pore perimeters $U_{\text{P}}$ were calculated with the *ImageJ Analyze Particles* function.

4. The mean apparent pore diameter $d_{\text{Apparent}}$ was determined for *n* pores by

$$d_{\text{Apparent}} = \frac{1}{n}\sum \frac{4A\text{p}}{U_{\text{p}}}. \tag{S4.2}$$

The above processing steps are illustrated in Fig. S4.1.

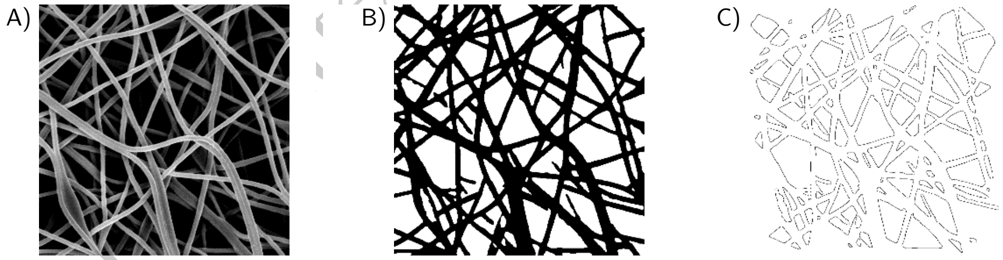


**Fig. S4.1:** Illustration of the SEM image processing steps for the apparent pore diameter and apparent porosity calculations. A) Threshold adjusted image with 6 to 7 visible fiber layers. B) Segmented image. C) Pore outlines.

Preprint [illegible]ed to ChemElectroChem

##### S4.2.2 3D macropore model

The 3D macropore model builds upon the linear relation between the macropore diameter and the fiber diameter (Eq. S4.4) derived in the previous section. It relies on the assumption, that the true three-dimensional structure of the electrospun materials does not yield a substantially different pore structure as the model. The deviations of the model from the true material morphology and their expected impact on the model accuracy are as follows:

1. The fiber diameters of the electrospun materials follow log-normal distributions [7]. The impact on the model accuracy is small, as the number of fibers per volume $n$ for a given porosity $\varphi$ is proportional to $(1-\varphi)/d_\mathrm{F}^2$ and the expected values of $\langle 1/d_\mathrm{F}^2 \rangle$ differ less than 9 % from $1/d_\mathrm{F,\,mean}^2$.

2. The fibers are bent and mostly not merged at fiber intersections. Both features have an opposing effect on the number of fibers per layer and compensate each other.

3. The fiber cross section is not perfectly circular. The impact on the number of fibers per layer can be considered small if the non-circular fiber cross sections are oriented randomly.

4. Interlaced fibers occur only occasionally due to the layer-by-layer fiber deposition during electro-spinnning and is therefore neglected. Furthermore, Nakamura et al. [8] found that the effect of fibers tilted against the *xy*-plane on the pore size distribution is negligible.

##### S4.2.3 Model construction

Randomly aligned fibers were constructed in the *xy*-plane with 800x800 $\mathrm{px}^2$ resolution. This resolution is sufficient to accommodate enough fibers for the apparent pore size calculation in a single layer. The fiber cross section was approximated as depicted in Fig. S4.2 B) and the corresponding fiber diameter calculated, assuming equal $A_\mathrm{Pixel}$ and $\mathrm{A_{Circle}}$, by $d_\mathrm{F} = \sqrt{4 \cdot A_\mathrm{Pixel}/\pi} = \sqrt{4 \cdot 13/\pi}\,\mathrm{px}$. Hundred fiber layers (corresponding to 500 px in the *z*-direction were stacked to form a 3D volume representation with a porosity of 95 %, similar to the porosity of the electropun materials. A 3D rendering of a representative sub volume is depicted in S4.2 A).

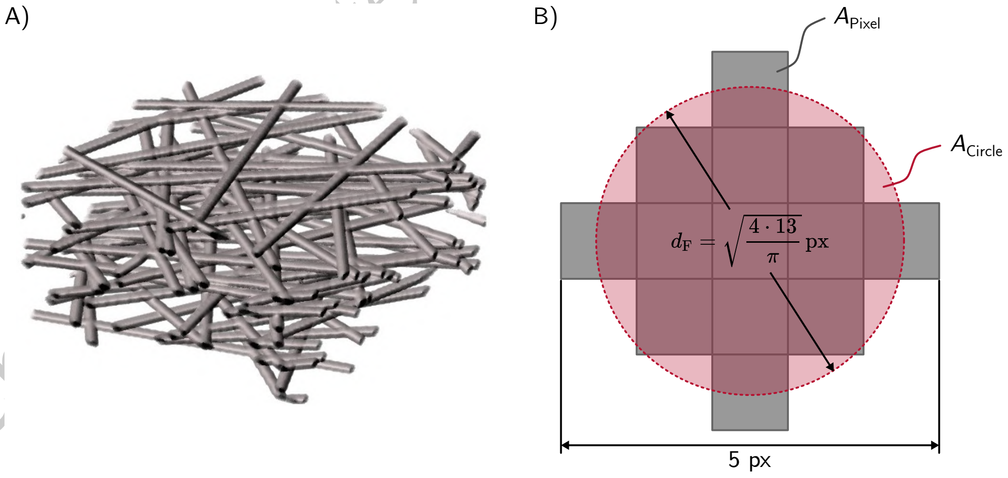


**Fig. S4.2:** A) 3D rendering of the fiber model. B) Pixel approximation of the circular fiber cross section.

##### S4.2.4 Apparent pore diameter and porosity

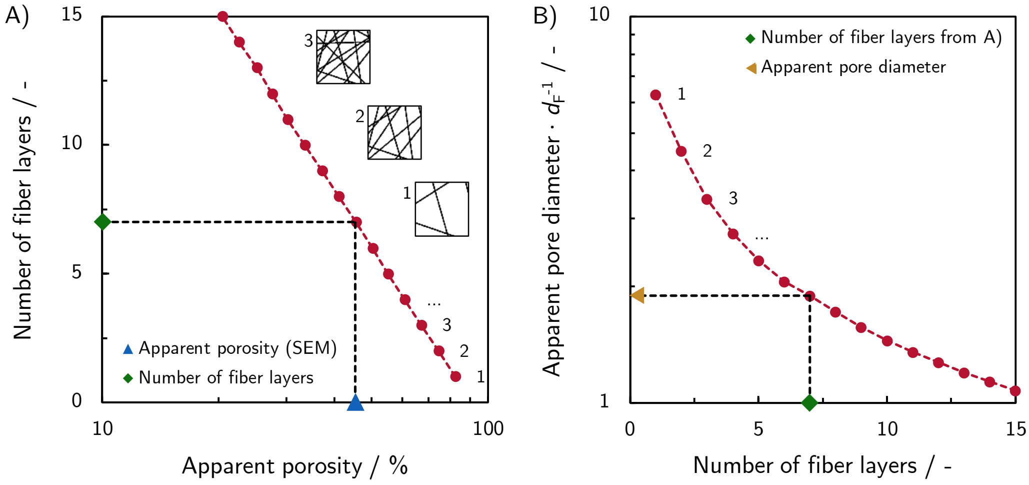


**Fig. S4.3:** Simulation of the apparent porosity and pore diameter. A) The apparent porosity allows the estimation of the number of fiber layers captured on the SEM image. B) The number of fiber layers translates in a slope for the prediction of the apparent pore diameter for a given fiber diameter according to Eq. S4.3.

The apparent porosity and apparent pore diameter of $z$-projections were analyzed according to the processing steps described in Section S4.2.1. Illustrations of the first 3 $z$-projections are given as inset in Fig. S4.3 A). The relation of the number of fiber layers and the apparent porosity $\varphi_{\text{Apparent}}$ depicted in Fig. S4.3 A) can be used to estimate the number of fiber layers that were captured in a SEM image as illustrated. The number of fiber layers translates into an apparent pore diameter normalized to the fiber diameter as illustrated in S4.3 B). This value represents a coefficient that relates the apparent pore diameter with the fiber diameter:

$$d_{\text{Apparent}} = \underbrace{\frac{d_{\text{Apparent}}}{d_{\text{F}}}(\varphi_{\text{Apparent}})}_{\substack{\text{Slope of model prediction} \\ \text{(Fig. S4.4)}}} \times d_{\text{F}} \qquad \text{(S4.3)}$$

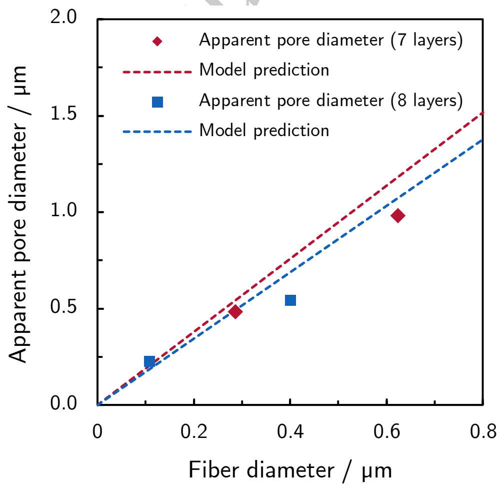


**Fig. S4.4:** Comparison of the apparent pore diameters mesured by SEM image analysis and the model predictions. The model deviates by less than 22 % from the values determined for the electrospun materials by SEM image analysis.

Pre... umitted to ChemElect... em

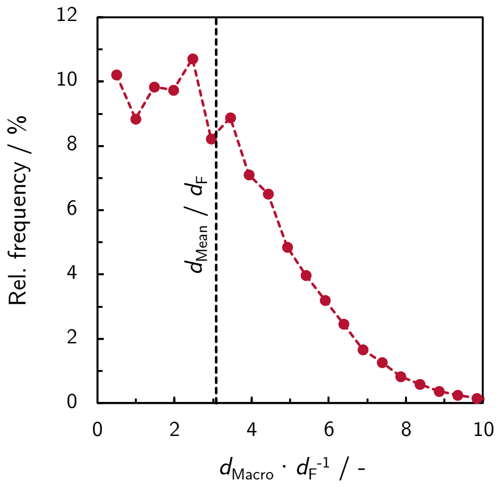


**Fig. S4.5:** Macropore diameter distribution normalized to the fiber diameter $d_F$ derived from the continuous pore size distribution. $d_{Mean}/d_F$ translates into the slope of the model prediction in Fig. S4.6.

##### S4.2.5 3D continuous pore diameter distribution

Using the *ImageJ Xlib* plugin [9] the 3D continuous pore diameter distribution of the 3D fiber model was calculated. The plugin fills the pore space with spheres of different diameters. The resulting pore diameter distribution is depicted in S4.5. From the pore diameter distribution the mean pore diameter normalized to the fiber diameter $d_{Mean}$ / $d_F$ was calculated.

##### S4.2.6 Conversion of apparent pore diameter to 3D pore diameter

The macropore diameter $d_{Macro}$ can be calculated from the model predictions (marked with an asterisk), the apparent pore diameter $d_{Apparent}$, and the apparent porosity $\varphi_{Apparent}$ determined from SEM images as follows:

$$d_{Macro} = \underbrace{\frac{d^*_{Mean}}{d^*_{Apparent}(\varphi_{Apparent})}}_{\text{Conversion factor}} \times d_{Apparent} \tag{S4.4}$$

The conversion factor depends on the number of fiber layers that are captured by the SEM image. The number of captured fiber layers can be estimated with the apparent porosity and the apparent pore diameter according $d^*_{Apparent}$ to Fig. S4.3. The macropore diameters of the electrospun fiber materials converted from SEM image analysis are depicted in Fig. S4.6 together with the model prediction. In summary: The 3D model predicts accurately the apparent porosity and the apparent pore diameter for a given fiber diameter. A conversion factor from the apparent macropore diameter, measured by analysis of the SEM images, to the 3-dimensional spherical pore diameter can be deduced from the 3D simulation.

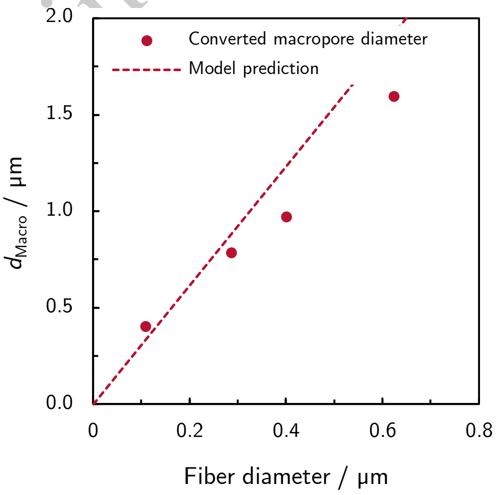


**Fig. S4.6:** Comparison of the macropore diameter model with according to Eq. S4.4 converted values. The slope of the model prediction is $d_{Mean}/d_F$. The deviation of the model prediction to the experimentally determined values is 20 %.

Preprint sumitted to ChemElectroChem

#### S5 Literature analysis

| | | | | | | |
|---|---|---|---|---|---|---|
| | p < .001 | | | | | |
| | Materials with mean pore diameter above 26 µm | | | Woven and knitted materials with broad pore size distribution, including small macropores | | |
| | TP-120 | GFD 2 | V1-Felt | C-Tex 27 | C-Tex 13 | C-Tex 20 |
| | 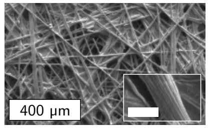 | 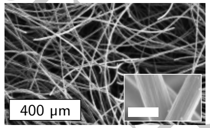 | 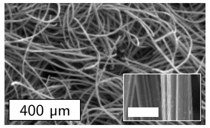 | 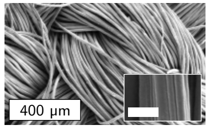 | 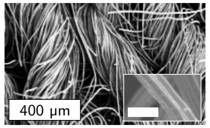 | 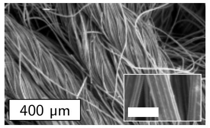 |
| Fiber diameter / µm | ∼ 8.5 | ∼ 8 | ∼ 10 | ∼ 16 | ∼ 7 | ∼ 7 |
| Fiber cross section | Grooved | Smooth | Grooved | Grooved | Grooved | Grooved |
| Fiber alignment | Random in plane | Random | Random | Dense, woven Aligned bundles | Loose, knitted Twisted bundles | Loose, knitted Twisted bundles |
| Thickness / mm | 0.37 | 2 | 1.8 | 0.5 | 0.5 | 1 |
| $d_{\mathrm{Macro}}$ / µm | 26[b] | 31 | 28 | - | - | - |
| Porosity[a] / % | 74 | 95 | 96 | 85 | 81 | 86 |
| $V_{\mathrm{Meso}}$ / $10^{-3}$ cm$^3$ | 0.4 | 2.0 | 4.8 | 2.3 | 3.4 | 5.5 |
| $V_{\mathrm{Micro}}$ / $10^{-6}$ cm$^3$ | 0.2 | 0.4 | 15.2 | 11.9 | 15.3 | 22.0 |
| $SRF_{\mathrm{BET}}$ | 1690 | 10920 | 161660 | 103950 | 132000 | 166320 |
| Current density / µA cm$^{-2}$ | 12 ± 2 | 12.0 ± 0.2 | 14.0 ±1.0 | 18.0 ± 0.4 | 24.0 ± 0.3 | 29 ± 0.3 |

**Table S5.3:** Overview of the material properties investigated by Kipf et al. [10] including additional morphological characteristics. The porosity was calculated assuming a carbon density of 1.75 g cm$^{-3}$. The current production was recorded in a 3-electrode configuration using a quasi-galvanostatic load curve and evaluated at -200 mV vs SCE. We found a significant higher current production for the knitted carbon cloths (featuring small macropores), compared to the materials with the materials that feature macropores with mean diameters above 26 µm ($p < .001$, Welch corrected one tail $t$-test). [a] The porosity has been calculated based on the material thickness provided by the manufacturers and have not been verified by own measurements, except for GFD 2. [b] The value may only serve as rough estimate due to binder residues and the low porosity. The scale bars in the insets correspond to 10 µm. Images reprinted from *Bioresource Technology*, 146, Elena Kipf, Julia Koch, Bettina Geiger, Johannes Erben, Katrin Richter, Johannes Gescher, Roland Zengerle, Sven Kerzenmacher, *Systematic screening of carbon-based anode materials for microbial fuel cells with Shewanella oneidensis MR-1*, pp 386–392, Copyright (2013), with permission from Elsevier.

Preprint submitted to ChemElectroChem

| | PAN | PAN-AC | PAN-GR |
|---|---|---|---|
| | 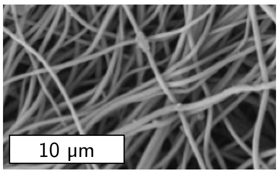  | 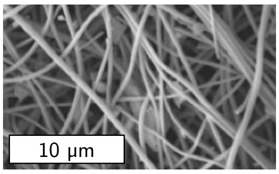  | 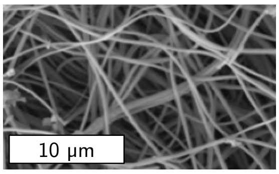  |
| Fiber diameter / nm | 434 ± 34 | 383 ± 79 | 327 ± 69 |
| Particles | none | Activated carbon | Graphite |
| $S_{\mathrm{BET}}$ / m$^2$ g$^{-1}$ | 168.6 | 310.0 | 362.1 |
| $V_{\mathrm{Micro}}$ / cm$^3$ g$^{-1}$ | 0.048 | 0.116 | 0.136 |
| $V_{\mathrm{Total}}$ / cm$^3$ g$^{-1}$ | 0.142 | 0.202 | 0.239 |
| $V_{\mathrm{Total}}$ - $V_{\mathrm{Micro}}$ / cm$^3$ g$^{-1}$ | 0.094 | 0.086 | 0.103 |
| Mean $j_{\mathrm{Max}}$ / µA cm$^{-2}$ | 114 ± 6.6 | 139 ± 8.5 | 155 ± 11.3 |

**Table S5.4:** Overview of the material properties investigated by Patil et al. [11]. The fibers were electrospun from 10 wt% PAN dissolved in N,N-dimethylformamide. The solution for the preparation of PAN-AC and PAN-GR contained additional 1 wt% activated carbon or graphite particles. The particles are visible between the fibers. The amount of spinning dope is 25 mL for all materials suggesting a similar area density of the materials (values not given in the publication). Images reprinted from *Bioresource Technology*, 132, Sunil A. Patil, Samuel Chigome, Cecilia Hägerhäll, Nelson Torto , Lo Gorton, *Electrospun carbon nanofibers from polyacrylonitrile blended with activated or graphitized carbonaceous materials for improving anodic bioelectrocatalysis*, pp 121–126, Copyright (2013), with permission from Elsevier.

Preprint sumitted to ChemElectroChem

| | CM | SB |
|---|---|---|
| | 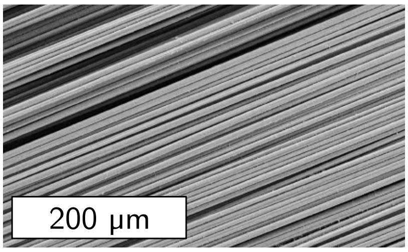  | 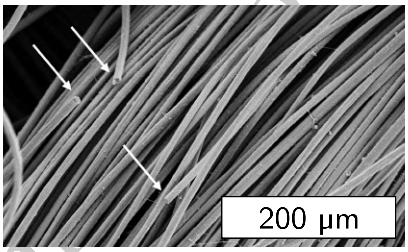  |
| Fiber diameter / µm | 7.5 | 10 |
| Fiber alignment | Dense bundles | Loose bundles |
| Carbon content | Maximum current density $j_{\text{Max}}$ / µA cm$^{-2}$ | |
| Low | $\sim 128 \pm 22$ | $\sim 92 \pm 34$ |
| Moderate | $\sim 85 \pm 10$ | $\sim 187 \pm 58$ |
| High | $\sim 117 \pm 17$ | $\sim 167 \pm 52$ |
| Mean $j_{\text{Max}}$ / µA cm$^{-2}$ | $110 \pm 22$ | $148 \pm 50$ |

**Table S5.5:** Overview of the material properties investigated by Pötschke et al. [12] including additional morphological characteristics and the corresponding current densities. The current densities were manually extracted from from bar charts. The current production was recorded in a 3-electrode configuration at 200 mV vs Ag/AgCl$_{\text{Sat}}$. We did not find a significant difference of the the mean current production between CM and SB materials ($p = .32$, Welch corrected $t$-test).
Images reprinted from *Frontiers in Energy Research*, 7 (2019), article 100, Liesa Pötschke, Philipp Huber, Sascha Schriever, Valentina Rizzotto, Thomas Gries, Lars M. Blank, Miriam A. Rosenbaum, *Rational Selection of Carbon Fiber Properties for High-Performance Textile Electrodes in Bioelectrochemical Systems*.

Preprint submitted to Chem[illegible]Chem


Preprint submitted to ChemElectroChem
